# Brain implantation of tissue-level-soft bioelectronics via embryonic development

**DOI:** 10.1101/2024.05.29.596533

**Authors:** Hao Sheng, Ren Liu, Qiang Li, Zuwan Lin, Yichun He, Thomas S. Blum, Hao Zhao, Xin Tang, Wenbo Wang, Lishuai Jin, Zheliang Wang, Emma Hsiao, Paul Le Floch, Hao Shen, Ariel J. Lee, Rachael Alice Jonas-Closs, James Briggs, Siyi Liu, Daniel Solomon, Xiao Wang, Nanshu Lu, Jia Liu

## Abstract

The design of bioelectronics capable of stably tracking brain-wide, single-cell, and millisecond-resolved neural activities in the developing brain is critical to the study of neuroscience and neurodevelopmental disorders. During development, the three-dimensional (3D) structure of the vertebrate brain arises from a 2D neural plate^1,2^. These large morphological changes previously posed a challenge for implantable bioelectronics to track neural activity throughout brain development^3–9^. Here, we present a tissue-level-soft, sub-micrometer-thick, stretchable mesh microelectrode array capable of integrating into the embryonic neural plate of vertebrates by leveraging the 2D-to-3D reconfiguration process of the tissue itself. Driven by the expansion and folding processes of organogenesis, the stretchable mesh electrode array deforms, stretches, and distributes throughout the entire brain, fully integrating into the 3D tissue structure. Immunostaining, gene expression analysis, and behavioral testing show no discernible impact on brain development or function. The embedded electrode array enables long-term, stable, brain-wide, single-unit-single-spike-resolved electrical mapping throughout brain development, illustrating how neural electrical activities and population dynamics emerge and evolve during brain development.

## Main text

Recording neural activities throughout brain development is critical to understanding how neurons self-assemble into an organ capable of learning, behavior, and cognition^1^. To date, however, it has not been possible to conduct the ideal experiment: recording brain-wide neural activity at the cellular level with millisecond temporal resolution in animals throughout brain development^3–6^. For example, functional magnetic resonance imaging (fMRI) acquires brain-wide activity non-invasively^7^ but has low spatiotemporal resolution. Microscopic imaging of genetically encoded calcium or voltage reporters can enable tissue-wide, cell-specific imaging. Using such tools, the brain-wide calcium activity of a few transparent species (e.g., larval zebrafish) has been studied^8^. Tissue scattering and 3D volumetric scanning, however, limit fast optical access to cells throughout 3D brains in non-transparent animals.

Implanted microelectrodes have the potential to overcome these limitations. Existing technologies support the interrogation of single-unit activity with millisecond temporal resolution in 3D tissue. Multielectrode arrays can simultaneously track electrical activities from a large number of neurons in the brain without sacrificing spatial or temporal resolution^10–13^. The developing brain, however, presents additional challenges. Vertebrate brains are complex 3D structures that originate from a 2D single-cell layer in the embryo. During development, cell division, proliferation, and folding events result in rapid volume expansion and large changes in tissue morphology^14–16^. Conventional microelectronics cannot interrogate such dynamically changing environments with accuracy, compatibility, and longevity due to fundamental mechanical mismatches between electronic and biological materials.

Recent advances in flexible electronics^17–21^ have yielded “tissue-like” mesh microelectronics: sub-micrometer-thick mesh electronics with tissue-level flexibility. Mesh microelectronics mimic the physicochemical properties of the extracellular matrix and are thus capable of seamlessly integrating into *in vitro* and *in vivo* neural tissues, forming gliosis-free bioelectronic interfaces and enabling long-term, stable, multichannel electrical recordings in 3D brain tissues at single-unit, single-spike spatiotemporal resolution over months^22^. Neurons, however, are innervated at the nanometer scale. Regardless of how small and soft bioelectronics are designed to be, the implantation of such devices into the mature brain necessarily introduces acute damage to neural tissue. We previously showed that stretchable electronics can be integrated into developing biological tissue *in vitro* via normal developmental processes^23,24^. The forces exerted by the developing tissue unfold and reconfigure the bioelectronics, distributing sensors throughout the 3D structure.

Here, we report the development and validation of a class of bioelectronic devices and methods to integrate the device within the developing brain of vertebrate animals throughout embryogenesis. Specifically, we have developed sub-micrometer-thick, tissue-level soft mesh electronics containing a stretchable electrode array that can be implanted into the embryo neural plate. During organogenesis, the neural plate undergoes a 2D-to-3D reorganization process^2^, folding, proliferating, and expanding into the precursors of the nervous system. The endogenous forces involved in this process seamlessly and non-invasively distribute and integrate the sensor network across the 3D volume of the neural tube and brain, creating a “cyborg” embryo. (We will refer to embryos and tadpoles which have embedded electronics as cyborg embryos and cyborg tadpoles going forward. Control tadpoles and embryos contain no embedded electronics.) The study revealed that the presence of microelectronics had no discernible impact on embryo development or subsequent behaviors. Our device enables tissue-wide, continuous recording of neuron electrical activity at millisecond temporal resolution. We validate our device by using it to illustrate the emergence of electrical activities in the neural plate and the evolution, synchronization, and propagation of neural dynamics over the time course of embryonic brain development.

### Soft and stretchable bioelectronics for minimally invasive brain implantation via embryonic neurulation

During vertebrate development, the neural plate, a 2D single-cell ectoderm-derived layer on the surface of the embryo, folds to form the neural tube, and, with further expansion and more folding, morphs into the 3D brain and other portions of the nervous system^2^. We hypothesized that this 2D-to-3D reconfiguration process could be leveraged to distribute appropriately designed soft, stretchable bioelectronics throughout the brain with minimal impact on brain development or function. We initially sought to test this hypothesis in the *Xenopus laevis* (frog) embryo (Fig. 1a), given that it is a widely used in developmental biology and because its developmental processes are well understood^1^. Moreover, the neural plate is exposed to aqueous solution during *Xenopus* embryogenesis^1^, making it accessible for the placement of a bioelectronic device prior to the neural tube formation. Figure 1b and 1c show two schematic views of how neural development drives the integration of an appropriately designed device. Mesh electronics containing a stretchable electrode array are implanted non-invasively, driven by the 2D-to-3D reconfiguration of neural tissue during neurulation (Fig. 1b). As the neural tube forms into a 3D structure, the stretchable electrode array fully integrates with neural networks throughout the entire brain (Fig. 1c), revealing the brain-wide electrophysiological evolution over the course of development (Fig. 1a).

**Fig. 1.**
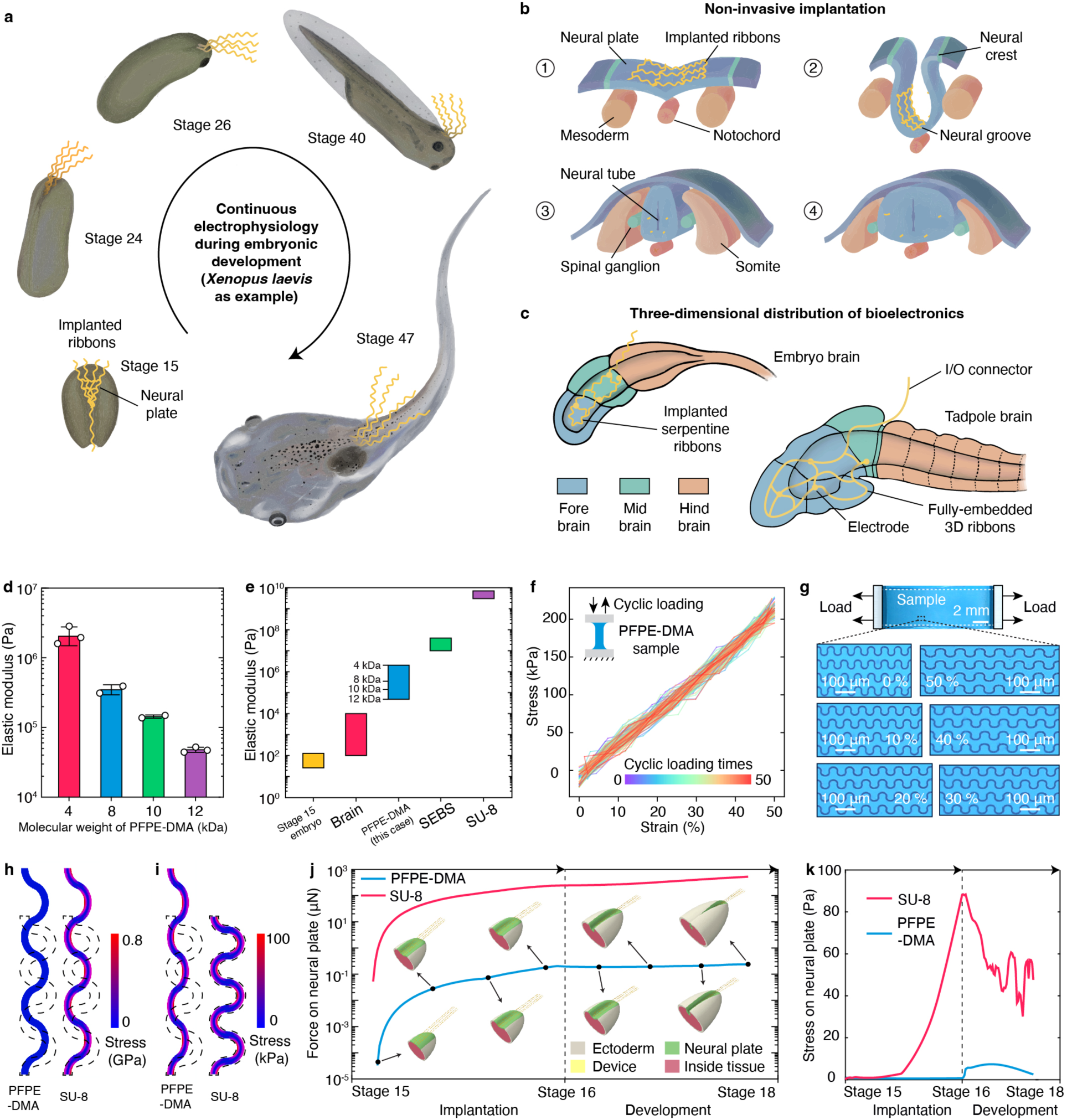
Design of soft and stretchable bioelectronics for brain implantation via embryonic development. **a,** Schematics showing the stepwise implantation of soft and stretchable mesh electronics into the brain of the *Xenopus* embryo via organogenesis. The mesh electronics track brain-wide, continuous electrophysiological evolution throughout brain development. The mesh electronics with a stretchable electrode array are laminated onto the neural plate at the beginning of neurulation (stage 15). During neurulation, tissue reconfiguration embeds the mesh into the neural tube (stage 24). The mesh deforms with the neural tube as it re-organizes into a 3D tadpole brain (stages 26, 40, 47). **b,** Schematics of zoomed-in sagittal sections of neural plate development showing how the mesh electronics integrate non-invasively into the neural tube via neurulation. **c,** Schematics showing that expansion and folding of the neural tube fully distribute stretchable mesh electronics throughout the 3D structure of the brain. **d,** Elastic modulus of crosslinked perfluoropolyether-dimethacrylate (PFPE-DMA) with 4-, 8-, 10-, 12-kDa molecular weight. **e,** Elastic modulus of stage 15 embryos, brain tissue, PFPE-DMA, Styrene-Ethylene-Butylene-Styrene (SEBS), and SU-8. The dashed lines in the PFPE-DMA column indicate the average elastic modulus of PFPE-DMA with 4-, 8-, 10-, and 12-kDa molecular weights. **f,** Stress-strain curve of a representative 8 kDa PFPE-DMA film under 50 cycles of 50% uniaxial stretch loading. **g,** Photographic image of representative 8 kDa PFPE-DMA-encapsulated serpentine gold (Au) ribbons in a uniaxial stretch test. Zoomed-in views show the serpentine ribbons at 0%, 10%, 20%, 30%, 40% and 50% strain states. **h, i,** Stress distributions in PFPE-DMA and SU-8 serpentine ribbons when stretched to the same strain (**h**) and the same maximum von Mises stress (**i**). Dashed lines show the initial shapes of the ribbons. **j, k,** Simulations showing the force (**j**) and maximum von Mises strain (**k**) from the SU-8 and PFPE-DMA meshes applied to the neural plate during implantation and development from stages 15 to 18.

We aimed to restrict device implantation to the cranial neural plate, as this region ultimately forms the brain (the caudal neural plate extends axially to form the spinal cord). According to our calculations, the formation of the frog embryo brain should introduce less than 30% axial strain to the device over the course of development. Therefore, our initial design featured 40-nm-thick chromium/gold serpentine interconnects and an 800-nm-thick SU-8 encapsulation layer, an approach that can tolerate up to 30% strain in developing organoids *in vitro*^23^. However, the elastic modulus of frog embryos at the neurulation stage is significantly lower than that of organoids (Extended Data Fig. 1a); the SU-8 device was too stiff, cutting through the neural plate shortly after integration and critically damaging the embryo (Extended Data Fig. 1b,c).

The bending stiffness of each device layer can be calculated by the expression *Ebh*^3^, where *E*, *b*, and *h* are the elastic modulus, width, and layer thickness, respectively. The SU-8 encapsulation layer dominated overall device flexibility, largely due to its thickness and intrinsic stiffness. We therefore hypothesized that a stretchable device with softer dielectric layers would be more flexible and less likely to damage the embryonic tissue. We designed and tested a 1-μm-thick mesh structure made of a softer elastomer: styrene-ethylene-butylene-styrene (SEBS). This mesh was successfully embedded into the neural tube during neurulation without obvious damage to the embryo (Extended Data Fig. 1d). SEBS, however, is incompatible with sub-micrometer multilayer photolithography, which is necessary for the fabrication of stretchable mesh electronics.

We therefore developed a perfluoropolyether-dimethacrylate (PFPE-DMA)-based photoresist, which possesses both low elastic modulus and intrinsic stretchability (Extended Data Fig. 1e,f) as well as the chemical orthogonality needed for fabrication. By tuning the molecular weight of the PFPE-DMA precursor, we could alter the elastic modulus of the photopatterned PFPE-DMA film (Fig. 1d). More flexible PFPE-DMA films better matched the mechanical properties of brain tissue but had lower fabrication yield. We ultimately chose PFPE-DMA with a molecular weight of 8 kDa and an elastic modulus of ∼0.3 MPa (Extended Data Fig. 1e, see Supplementary discussion). For comparison, the elastic modulus of the stage 15 embryo and brain tissue are 74.10 ± 2.45 Pa (mean ± s.e.m., *n* = 100) and ∼1 kPa, respectively; while the elastic modulus of SU-8 and SEBS are ∼4 GPa^25^ and ∼17.65 MPa^26^, respectively (Fig. 1e and Extended Data Fig. 1a). In hysteresis testing, 8 kDa PFPE-DMA film maintained stable linear elasticity through 50 cycles of 50% uniaxial stretching (Fig. 1f). Finally, 8 kDa PFPE-DMA-encapsulated gold serpentine ribbons maintained their structure under 50% uniaxial stretching (Fig. 1g).

To further evaluate the potential impact of device stiffness on embryonic brain implantation, we compared PFPE-DMA and SU-8 encapsulated serpentine ribbons using finite element analysis (FEA). FEA showed that when stretching 1-μm-thick PFPE-DMA and SU-8 ribbons encapsulating 40-nm-thick gold to a 29% strain, close to the relative extension of the *Xenopus* brain during early-stage development, the maximum stress in PFPE-DMA was only 68 kPa while the maximum stress in SU-8 ribbon reached 759 MPa (Fig. 1h). As the yield strength of PFPE-DMA is 0.45 MPa (>> 68 kPa, Extended Data Fig. 1e), and the yield strength of SU-8 is 80 MPa^25^ (<< 759 MPa), the PFPE-DMA ribbon remained in the elastic deformation regime, while the SU-8 ribbon would likely fail. When the two ribbons were stretched to the same maximum stress of 100 kPa, close to the limit that the embryonic tissue can sustain^27^, the PFPE-DMA ribbon reached a 39.48% strain (Fig. 1i), exceeding embryonic axial deformation during development. In contrast, the SU-8 ribbon barely stretched at all (0.0040% strain), indicating that it is too stiff to co-develop with the embryo. FEA of the interaction between each mesh and the embryonic neural plate during neurulation (Fig. 1j,k and Extended Data Fig. 2, see Methods) showed that the 1-μm-thick PFPE-DMA mesh encapsulating 40-nm-thick gold introduced a maximum force of ∼0.1 μN and maximum stress of ∼10 Pa to the neural plate. The SU-8 mesh introduced a maximum ∼1 mN of force and ∼90 Pa of stress. Thus, FEA suggested that mechanical deformation or damage to the neural plate during implantation resulting from PFPE-DMA meshes would likely be much lower than those caused by SU-8 meshes.

In addition to its mechanical compatibility, PFPE-DMA shows superior contact properties with embryos compared to SU-8. In implantation tests, SU-8 devices adhered to embryo cells, scraping cells from the embryos (Extended Data Fig. 1b). In contrast, PFPE-DMA devices had a lower surface free energy (21.2 mJ/m² compared to 29.5 mJ/m² for SU-8, Extended Data Fig. 1g) and a smaller contact angle with cytomembrane analog (23.66° ± 1.24° compared to approximately 0° for SU-8, mean ± s.e.m., n = 5, Extended Data Fig. 1h). Experimental testing supported these measurements: a 1-μm-thick PFPE-DMA stretchable mesh can be successfully implanted into the embryonic brain with no obvious damage (Extended Data Fig. 1i).

We next asked what device structure would minimize impact on embryonic development when integrated via our proposed method (Fig. 2a). First, to accommodate the neural plate’s folding, the device needed to maintain close contact with neural plate cells, ensuring it becomes embedded into the neural tube as it forms. The interconnects of the device should be deformable to accommodate the 3D re-organization of the neural tube and eventually enable device integration into the 3D structure of the tadpole brain. Therefore, the device should contain: (i) polymeric thin-film blockers to keep the stretchable mesh electrodes away from the caudal neural plate (which elongates to form the spinal cord) (Fig. 2a and Extended Data Fig. 1j,k); (ii) stretchable ribbons anchored on the substrate to hold the electrodes at the correct portions of the neural plate (Fig. 2a and Extended Data Fig. 1k,l); and (iii) face-down electrodes connected via fully encapsulated stretchable interconnects (Supplementary Fig. 1a, see Supplementary discussion).

**Fig. 2.**
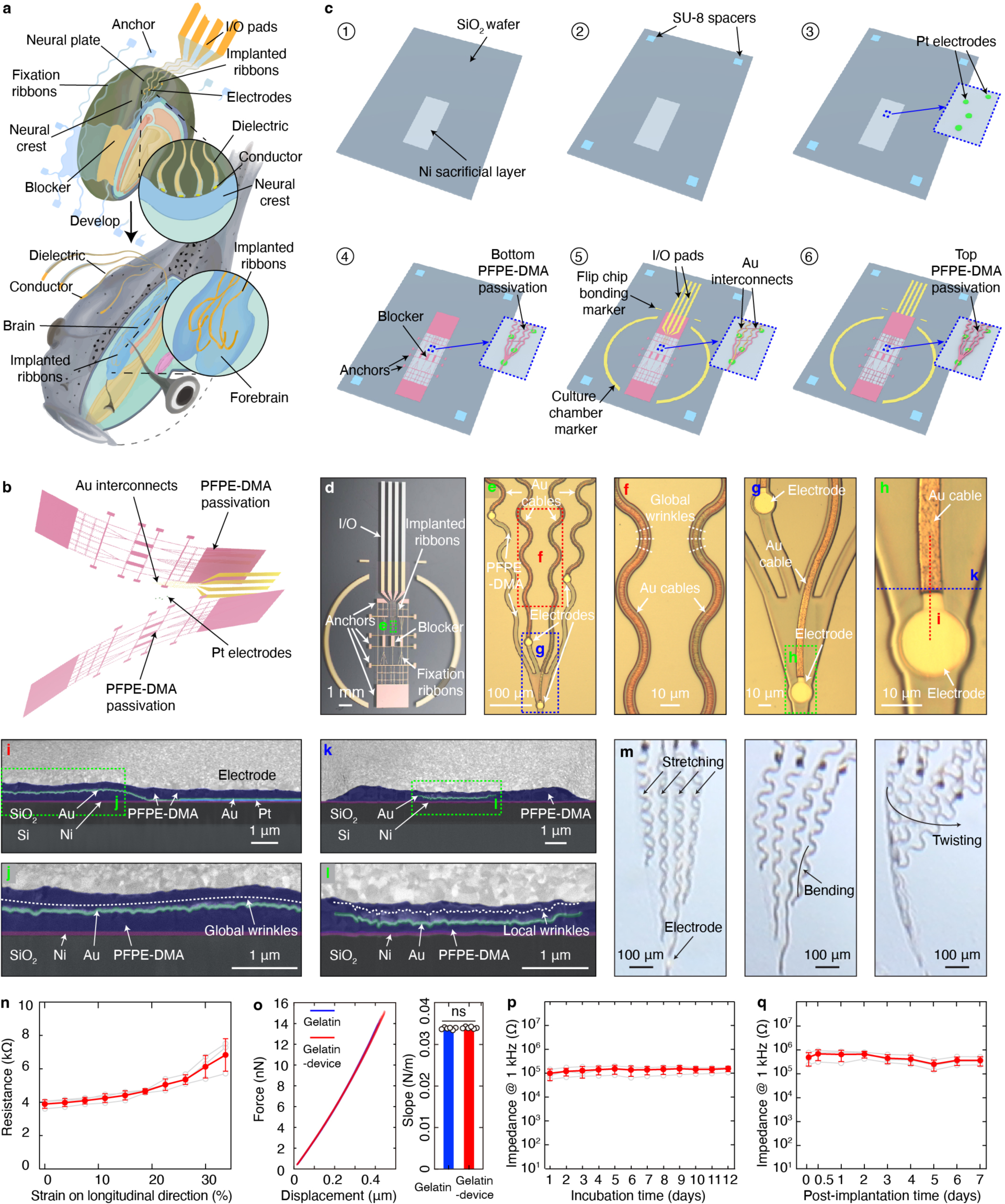
Fabrication of tissue-level-soft stretchable mesh electronics for brain implantation via embryo development. **a,** Schematics illustrating the design of soft and stretchable mesh electronics for embryo implantation. The design includes a stretchable mesh electrode array for electrophysiological sensing, interconnects and input/output (I/O) pads for data collection, polymeric stretchable anchors to hold the mesh to the neural plate, and blockers to restrict the mesh to the cranial neural plate. **b,** Schematic showing the tri-layer structure of the PFPE-DMA-encapsulated stretchable mesh electronics: PFPE-DMA passivation layers sandwich the Au interconnects layer. The electrodes are made of platinum (Pt) and electroplated with Pt-black. **c,** Fabrication flow of PFPE-DMA-encapsulated stretchable mesh electronics. First, a nickel (Ni) layer is deposited on a silicon oxide wafer as a sacrificial layer (step 1). A SU-8 layer is patterned as a spacer (step 2). Then, Pt electrodes are photolithographically patterned (step 3). Finally, the bottom PFPE-DMA layer (step 4), Au interconnects (step 5), and top PFPE-DMA layer (step 6) are photolithographically patterned. Insets highlighted by the blue dashed box show zoomed-in views of the stretchable mesh electrode array. **d,** Photographic image of representative PFPE-DMA mesh electronics on a glass substrate. **e,** Bright-field (BF) microscopic image of the zoomed-in view of the green dashed box-highlighted region in (**d**) showing the stretchable mesh electrode array for electrophysiological recording. **f,** BF microscopic image of the zoomed-in view of the red dashed box-highlighted region in (**e**) showing the stretchable interconnects. **g,** BF microscopic image of the zoomed-in view of the blue dashed box-highlighted region in (**e**), showing Pt electrodes. **h,** BF microscopic image of the green dashed box-highlighted region in (**g**) showing an individual electrode. **i-l,** Scanning electron microscope (SEM) images showing cross-sections of PFPE-DMA-encapsulated Au interconnects. Each layer is pseudo-colored and labeled. **i**, SEM image showing longitudinal cross-section along the red dashed line in (**h**). **j,** SEM image of the zoomed-in view of the green dashed box-highlighted region in (**i**). A dashed line is drawn parallel to the Au layer to indicate its global wrinkles. **k,** SEM image showing transverse cross-section along the blue dashed line in (**h**). **l,** SEM image of the zoomed-in view of the green dashed box-highlighted region in (**k**). A dashed line is drawn parallel to the Au layer to indicate its local wrinkles. **m,** Photographic images showing the free-floating stretchable mesh electrode array during stretching, bending, and twisting tests. **n,** Resistance as a function of strain during the longitudinal stretch test of PFPE-DMA-encapsulated electronics. Red dots and line plots indicate mean ± s.d., and each gray dot and line plot represents one sample. **o,** (Left) force as a function of displacement in atomic force microscope three-point bending tests of gelatin membrane and gelatin membrane embedded with PFPE-DMA mesh electronics showing tissue-level softness and stretchability of the device. (Right) statistics of curve slopes in the left figure. **p,** Electrode impedance at 1 kHz in 37 °C PBS as a function of incubation time. Red dots and line plots indicate mean ± s.d., and each gray dot and line plot represents one sample. **q,** Electrodes’ impedance at 1 kHz as a function of post-implantation time. Red dots and line plots indicate mean ± s.d., and each gray dot and line plot represents one sample.

### Fabrication of soft, stretchable mesh electronics

We sought to implement PFPE-DMA as the encapsulation layer in the fabrication of a functional stretchable mesh device. We adapted a typical microfabrication protocol to enable the photopatterning of multilayered ultra-thin PFPE-DMA structures as follows (Fig. 2b,c, see Methods): (i) As dimethacrylate polymerization is sensitive to oxygen, we contained the conventional mask aligner in a nitrogen chamber to prevent spin-coated PFPE-DMA films from oxygen exposure (Extended Data Fig. 3a-d); (ii) Photoresist and metal were initially difficult to deposit on PFPE-DMA films (Extended Data Fig. 3e). To improve surface adhesion, we treated the PFPE-DMA surface with argon gas plasma, allowing for photopatterning of standard photoresist and metal deposition on the PFPE-DMA layer to form interconnects (Extended Data Fig. 3f and Supplementary Fig. 1b); (iii) The platinum (Pt) electrode array was patterned as the bottom layer (Fig. 2b,c and Extended Data Fig. 3g) to enable direct contact with neurons during implantation.

We then evaluated the mechanical properties of the PFPE-DMA device. Photographic and bright-field (BF) microscopic images of the device (Fig. 2d-h) showed that the PFPE-DMA passivation and gold interconnects were successfully patterned with micrometer resolution. Scanning electron microscope (SEM) images (Extended Data Fig. 4a,b) and atomic force microscope (AFM) topographic images (Extended Data Fig. 4c,d) showed smooth PFPE-DMA passivation surfaces without discernible cracks or flaws. SEM images of the device cross-section (Fig. 2i-l) showed that the gold interconnects were fully encapsulated by the PFPE-DMA passivation. SEM, AFM, and BF microscopic images indicated that the PFPE-DMA-encapsulated gold interconnects self-wrinkled at multiple scales: anisotropic global wrinkles at the micrometer scale (Fig. 2j) perpendicular to the longitudinal direction of gold interconnects (Extended Data Fig. 4e) and isotropic local wrinkles at the sub-micrometer scale (Fig. 2l and Extended Data Fig. 4f,g). Together, the multi-scale wrinkles give the gold interconnects with a high degree of stretchability. After fabrication, the device was released from the substrate (Extended Data Fig. 3h). The free-floating stretchable mesh device showed no clear damage when stretched, bent, or twisted (Fig. 2m). After stretching and bending, SEM images revealed the device ribbons remained intact with passivation (Extended Data Fig. 4h-j). Impedance testing demonstrated that the fully encapsulated interconnects sustained conductivity after bending, 33% longitudinal or 38% transverse uniaxial strain (Fig. 2n and Extended Data Fig. 4k,l). In three-point bending test using AFM, the device introduced negligible additional force when embedded inside a 100-µm-thick gelatin membrane as compared to gelatin with no embedded device (Fig. 2o, see Methods), suggesting tissue-level softness and stretchability.

We then characterized the electrical performance and *in vitro* biocompatibility of stretchable electronics. Electrodes were electroplated with Pt-black to further reduce their electrochemical impedance for *in vivo* recording (Extended Data Fig. 4m). Electrode impedance was consistent from batch-to-batch (Extended Data Fig. 4n), indicating the robustness of sensor fabrication procedures. Continuous measurements of device electrochemical performance showed stable impedance over the 12-day time course of *in vitro* incubation (Fig. 2p) and over the 7-day time course of *in vivo* implantation (Fig. 2q). A 10-day *in vitro* co-culture of the PFPE-DMA or its potential degradation products with wild-type rat cortical neurons did not result in a significant change in the live/dead cell ratio of the neurons (*n* ≥ 6, *p* > 0.05; Extended Data Fig. 4o. All statistical tests performed were two-tailed, unpaired t-tests unless specified otherwise; actual p values are reported in Supplementary Table 1). Together, the longevity, stretchability and *in vitro* biocompatibility of the mesh electronics were sufficient to move forward with implantation and *in vivo* electrophysiological interrogation throughout *Xenopus* development.

### Brain-wide, non-invasive implantation of stretchable mesh electronics via embryo development

Next, we proceeded to implant the PFPE-DMA-based mesh electronics into the frog embryo at the Nieuwkoop and Faber stage 15. Time-lapse BF imaging (Fig. 3a) showed the gradual internalization of the stretchable mesh electronics by the neural plate as it bends to form the neural tube (Stages 15 to 19). Images from stages 30 to 47 (Fig. 3b-d) showed that as the embryo developed into a tadpole, the stretchable meshes were completely embedded into the brain without interrupting its development. A flexible cable connected the electrode array to the exterior of the brain for data collection. Once the devices were released from their substrates, cyborg frog tadpoles could swim freely with the embedded device. Cyborg frog tadpoles exhibited normal development up to later stages, showing no significant differences in survival ratio or developmental stages compared to the control group (Supplementary Fig. 2).

**Fig. 3.**
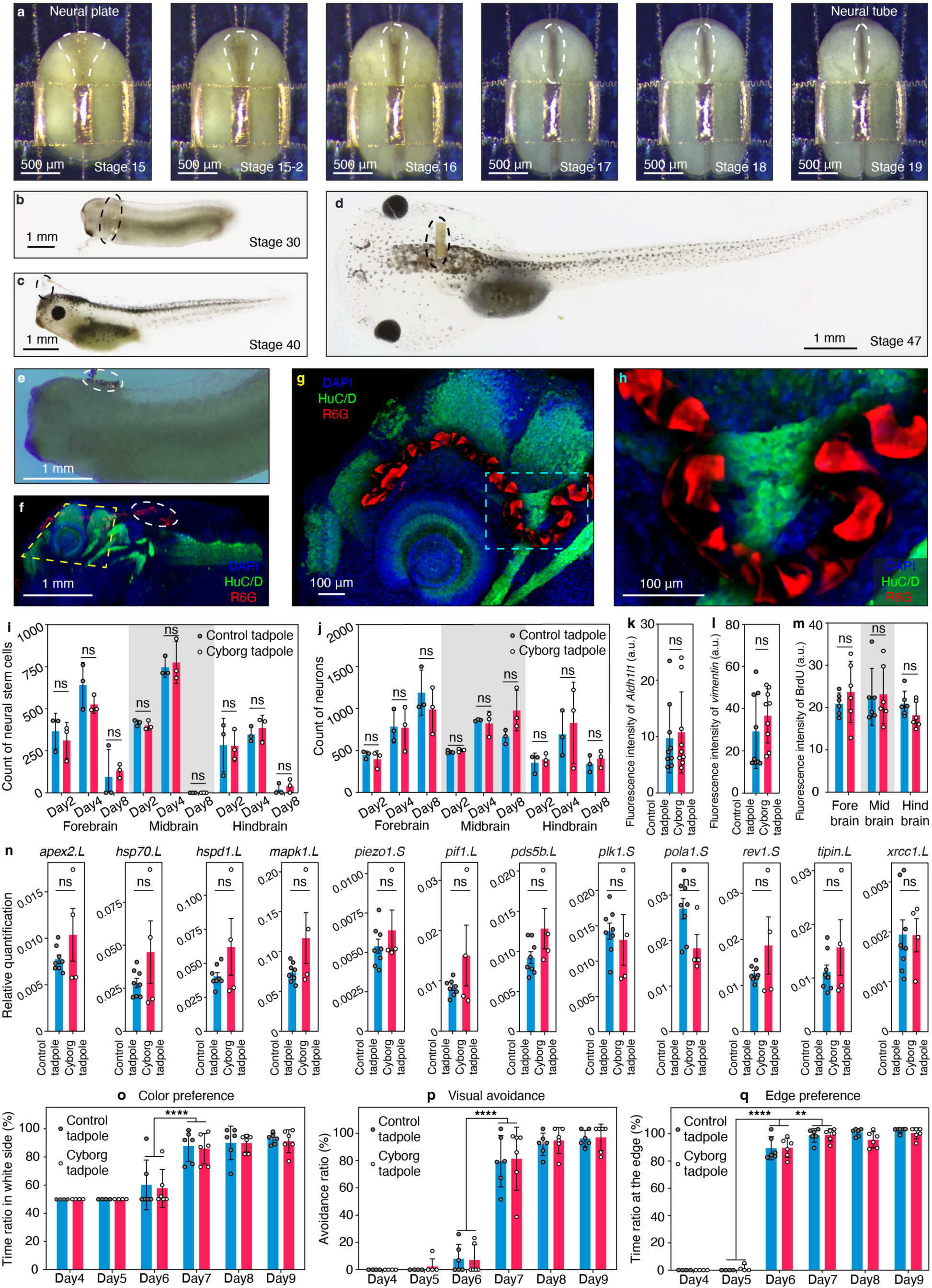
Minimally invasive brain implantation of tissue-level-soft, stretchable mesh electronics via embryonic development. **a,** Time-lapse BF microscopic images of a representative frog embryo implanted with stretchable mesh electronics at different development stages showing the gradual internalization of the mesh electrode array (dashed circles) into the neural plate. **b-d,** Optical photographic images of the embryo implanted with stretchable mesh electronics at stages 30 (**b**), 40 (**c**), and 47 (**d**). The dashed circles highlight the interconnects outside the brain. **e, f,** Photograph (**e**) and 3D reconstructed confocal fluorescence image (**f**) of a fixed cyborg tadpole. White dashed circles highlight the stretchable interconnects extending from the sensor array to the outside of the brain. 4′,6-diamidino-2-phenylindole (DAPI) labels cell nuclei, Rhodamine 6G (R6G) labels device, HuC/D labels neurons. **g,** Zoomed-in image of the yellow tilted dashed box-highlighted region in (**f**) showing the stretchable mesh electronics folded inside the neural tube. **h,** Zoomed-in image of the cyan dashed box-highlighted region in (**g**) showing the interface between the embedded stretchable mesh electronics and neurons. **i, j,** Bar and dot plots showing the number of neural stem cells (**i**) and neurons (**j**) identified from fluorescence images of the cryosection stained cyborg and control tadpoles at different developmental stages. **k-m,** Bar and dot plots showing the fluorescence intensity of aldehyde dehydrogenase 1 family member 1 (*Aldh1l1*) (**k**), *vimentin* (**l**), and bromodeoxyuridine (BrdU) (**m**) from fluorescence images of the cryosection stained cyborg and control tadpoles at stage 49. In **i-m,** Bar plots indicate mean ± s.d., each dot represents one sample, two-tailed unpaired t-test, ns, not significant. **n,** Quantitative polymerase chain reaction targeting stress genes of cyborg and control tadpoles. Bar and dot plots showing the stress gene expressions in cyborg and control tadpoles at stage 49. Bar plots indicate mean ± s.e.m., each dot represents one sample, two-tailed unpaired t-test, *n* ≥ 4, ns, not significant. **o-q,** Statistical analysis of color preference (**o**), visual avoidance (**p**), and edge preference (**q**) behavior test data from control and cyborg tadpoles at 4-to 9-days post fertilization. Bar plots indicate mean ± s.d., each dot represents a single trial. two-tailed unpaired t-test, *n* ≥ 8, **, *p* < 0.01, ****, *p* < 0.0001.

To characterize the 3D distribution of mesh electronics within the tadpole brain, frog tadpoles were fixed, cleared, whole-mount-stained, and imaged (details of antibody usage in this and subsequent staining experiments are provided in Supplementary Table 2). This approach ensures that the brain-electronics system remained intact throughout imaging (Extended Data Fig. 5a). We fixed the tadpole (Fig. 3e), cleared the tissue by removing the pigment and lipids^28^, and stained cell nuclei with 4′,6-diamidino-2-phenylindole (DAPI), neurons with HuC/D, and electronics with Rhodamine 6G (R6G). 3D reconstructed confocal fluorescence microscopic imaging (Fig. 3f) showed that the mesh electronics embedded across multiple brain regions. Zoomed-in images (Fig. 3g,h, Extended Data Fig. 5c and Supplementary Fig. 3) further illustrated the integration of the mesh within the 3D structure of the neural tissue. The device appears embedded into the forebrain, midbrain, and hindbrain, and the electrodes form close contact with neurons in several places. When we implanted the device at a later stage of neurulation (stage 16), the device did not appear in the neural tube but only superior to it (Extended Data Fig. 5d), demonstrating the necessity of integrating the mesh with the neural plate at the beginning of neurulation. We further carried out cell-type-specific protein marker staining, imaging coronal cryosections of the fore-, mid-, and hindbrains of tadpoles fixed at 2-, 4-, and 8-days post fertilization (DPF) (Extended Data Fig. 5b). The basal body, neurons, and neural stem cells were stained with acetylated-tubulin, myelin transcription factor 1 (Myt1), and SRY-box transcription factor 2 (Sox2), respectively. Confocal fluorescence microscopic images further confirmed device integration into brain tissue (Extended Data Fig. 5e).

To assess the impact of the embedded devices on cell proliferation and differentiation, and any potential immune response during development, we compared the cell number (Supplementary Fig. 4a) and fluorescence intensity (Supplementary Fig. 4b) of cyborg frog tadpoles with control frog tadpoles from cryosection fluorescence images at multiple time points during development. Results showed no statistically significant difference in neural stem cell or neuron numbers between cyborg and control tadpoles (for each brain region of a specific day, *n* ≥ 3 tadpoles, *p* > 0.05; Fig. 3i,j). In addition, *aldehyde dehydrogenase 1 family member 1* (*Aldh1l1*)^29^, *vimentin*^29^ and bromodeoxyuridine (BrdU)^30^ staining that quantify astrocyte numbers and cell proliferation at the site of a lesion, showed no statistically significant difference between control and cyborg tadpoles (for each brain region, *n* ≥ 6 tadpoles, *p* > 0.05; Fig. 3k-m, see Methods). These results are consistent with previous reports using flexible microelectronics for *in vivo* implantation^18,20,22, 31–35^.

Moreover, we examined stress genes to understand the potential effects of embedded electronics on embryo development. To conduct this analysis, we selected stress genes of *Xenopus laevis* from Xenbase^36^, with a specific emphasis on genes that have been extensively researched and are supported by a substantial number of references. Subsequently, we performed quantitative polymerase chain reaction (qPCR) on both cyborg and control embryos at 8-DPF. Our findings revealed no significant differences in the expression levels of the stress genes we assessed, including *apex2.L*, *hsp70.L*, *hspd1.L*, *mapk1.L*, *piezo1.S*, *pif1.L*, *pds5b.L*, *plk1.S*, *rev1.S*, *tipin.L* and *xrcc1.L* (Fig. 3n). These compelling results support that the presence of embedded microelectronics had minimal impact, if any, on the developmental processes of the embryos.

To further assess device integration impact on *Xenopus* development, we assessed frog tadpole behavior using three well-established behavioral tests (Extended Data Fig. 6a-i, see Methods): (i) color preference to characterize the visual function based on the preference of frog tadpoles to stay on the white side of a bicolored tank^37^; (ii) visual avoidance to characterize the maturation of visual responses in the optic tectum based on the ability of tadpoles to avoid incoming obstacles^38^; and (iii) edge preference to test the motor behavior of tadpoles based on their preference and ability to move along the edge of a container^39^. Analyses of behavior data showed that at 9-DPF, cyborg and control tadpoles (i) had no statistically significant difference in their preferences to stay in the white half of a bicolored tank (control vs. cyborg tadpoles, 92.8% vs. 91.2%, *n* ≥ 4 tadpoles, *p* > 0.05, Extended Data Fig. 6j); (ii) both avoided black dots introduced by a screen with no statistically significant difference in their success ratio (control vs. cyborg tadpoles, 100% vs. 97.2%, *n* ≥ 4 tadpoles, *p* > 0.05, Extended Data Fig. 6k); and (iii) swam along the edge of their container (the outmost quarter of the radius) at the same speed (∼0.4 mm/s) with no statistically significant difference in the proportion of time spent at the edge (control vs. cyborg tadpoles, 93.2% vs. 93.9%, *n* ≥ 4 tadpoles, *p* > 0.05, Extended Data Fig. 6l).

We further assessed these behaviors in control and cyborg tadpoles at 1- to 9-DPF on a daily basis. Data from 1- to 3-DPF were not included, as tadpoles still could not swim. Results showed no statistically significant difference between cyborg and control tadpoles at any stage of development (for each day, *n* = 6 tadpoles, *p* > 0.05; Fig. 3o-q). Notably, both cyborg and control tadpoles began to display the color preference at 7-DPF (from 6- to 7-DPF, *n* = 12 tadpoles, *p* < 0.0001; Fig. 3o), visual avoidance at 7-DPF (from 6- to 7-DPF, *n* = 12 tadpoles, *p* < 0.0001; Fig. 3p), and edge preference at 6-DPF (from 5- to 6-DPF, *n* ≥ 8, *p* < 0.0001; from 6- to 7-DPF, *n* = 12, *p* < 0.01; Fig. 3q). In total, behavioral assessment suggested that the implanted devices did not introduce a significant perturbation to the development of the tadpole visual and motor systems, nor did they impact tadpole visual and motor system function.

### Continuous tracking of neural electrophysiology over the course of *Xenopus* embryonic brain development

Subsequently, we tested whether the integrated electronics could enable continuous electrical recording from the developing brain and benchmarked signals with previous terminal measurements at various developmental stages. Cyborg tadpoles were cultured in an oxygen anesthetic system as previously reported to minimize tadpole movement during recording^40^ (Extended Data Fig. 7a). At this point, it was challenging to record from behaving tadpoles due to the restriction of tadpole motion by the flexible cable that connects the embedded electronics to the data acquisition system. The oxygen anesthetic system we adapted has been shown not to interrupt tadpole development^40^. We connected the mesh electronics via interconnects and input/output (I/O) pads to the amplification and data acquisition system using a flexible cable. Electrical recording was conducted in a Faraday cage (Extended Data Fig. 7b).

We continuously recorded electrical signals from the developing embryonic brains. Recordings from the same embryo at different timepoints showed the evolution of stage-specific electrical activities at different developmental stages (Fig. 4 and Extended Data Fig. 7c). In a representative experiment, at stage 24, we recorded spontaneous slow oscillations (amplitude: 941.0 ± 73.4 μV, width: 1.44 ± 0.07 s, interval: 3.26 ± 0.10 s, mean ± s.e.m., *n* = 3 tadpoles) from four electrodes distributed across different brain regions (Fig. 4a-c and Extended Data Fig. 7c-e). Analysis of the activation time delay across the channels (Fig. 4d,e, see Methods) showed that the oscillation waves maintained a constant propagation direction, from the forebrain to the midbrain, suggesting that brain-wide electrical activities were synchronized at this stage. Then, at stage 26, faster calcium wave-like signals emerged (Fig. 4f-h and Extended Data Fig. 8a,b, amplitude: 163.7 ± 5.2 μV, width: 132.3 ± 3.0 ms, interval: 1.9 ± 0.1 s, mean ± s.e.m., *n* = 3 tadpoles). The width and interval of the waveforms were consistent with recordings of spontaneous localized calcium release from cells reported previously^41^ (Extended Data Fig. 7f,g). Cross-channel examination of the temporal dynamics of these fast waves across different channels did not indicate a stable time delay, potentially indicating the gradual increase of localized activities in the brain (Fig. 4i,j, see Methods). Finally, at stage 40, both local field potential-like signals and fast spikes were observed in the brain (Fig. 4k-m). By applying a 300-3,000 Hz bandpass filter, we isolated single-unit action potential-like fast spikes using a conventional spike sorting algorithm (Fig. 4n, see Methods). The widths of the action potential (amplitude: 31.00 ± 0.69 μV, width: 2.04 ± 0.04 ms, interval: 41.83 ± 3.15 ms, mean ± s.e.m., *n* = 3 tadpoles) were comparable with previously reported data^42^ (Extended Data Fig. 7h). Drug testing ([2R]-amino-5-phosphonopentanoate [APV], cyanquixaline [CNQX], bicuculline [BIC], picrotoxin [PTX], and tetrodotoxin [TTX]) confirmed that these spikes came from the electrical activities of neurons (Extended Data Fig. 8c-g). After recording, we fixed, cryosectioned, and stained the tadpoles to identify electrode position in the brain (Extended Data Fig. 8h-o). Images showed that electrode which recorded single-unit action potential-like signals formed direct contacts with neurons in the brain (Extended Data Fig. 8l-o).

**Fig. 4.**
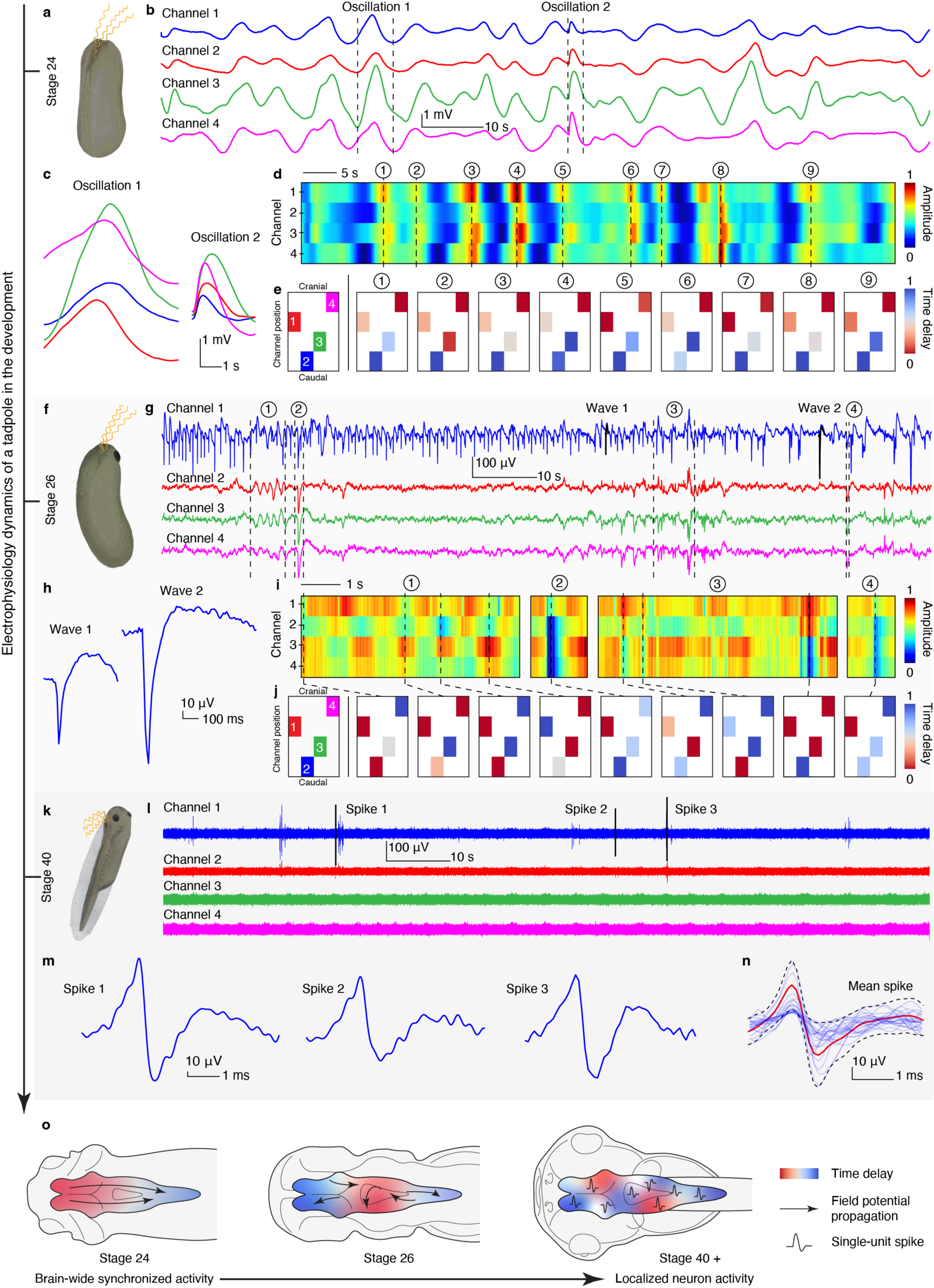
Continuous tracking of *in vivo* neural electrical activities from the same tadpole during organogenesis and brain development. **a,** Schematic of the cyborg tadpole at developmental stage 24. **b,** Representative voltage traces from four channels in the cyborg tadpole at stage 24. **c,** Zoomed-in views of the signals highlighted by dashed lines in (**b**). **d,** Heat map of the amplitude of the signals as a function of time for the voltage traces in (**b**). **e,** Spatiotemporal delay of signals across channels from the highlighted timepoints in (**d**). The figure on the left illustrates the positions of channels in the brain. **f,** Schematic of the cyborg tadpole at developmental stage 26. **g,** Representative voltage traces from four channels in the cyborg tadpole at stage 26. **h,** Zoomed-in views of the signals highlighted by dashed lines in (**g**). **i,** Heat map of the amplitude of the signals as a function of time for the voltage traces in (**g**). The figure on the left illustrates the positions of channels in the brain. **j,** Spatiotemporal delay of signals across channels from the highlighted timepoints in (**i**). **k,** Schematic of the cyborg tadpole at developmental stage 40. **l,** Voltage traces from four channels in the cyborg tadpole at stage 40. **m,** Zoomed-in views of representative single spikes highlighted by dashed lines in (**l**). **n,** Mean spike (mean ± s.d.) overlaid on all spikes from the same sorted unit, derived from the data presented in (**l**). **o,** Schematics illustrating how the neural activity evolves from brain-wide coordinated activity to localized neural activity and the emergence of single-unit spikes during *Xenopus* development.

### Tracking of single-unit action potential by high-density mesh electrode array during axolotl embryonic brain development

We next significantly increased the number and density of electrodes while maintaining the mechanical properties of the device to be compatible with the embryo development. This advancement allows the electrical activities from the same neurons to be simultaneously mapped by multiple electrodes, improving spike sorting accuracy and enabling the ability to track the neurons in developing tissue. To achieve this, we further explored electron-beam (e-beam) lithography to pattern nanometer-wide metal interconnects on the soft PFPE-DMA substrate (Extended Data Fig. 9a). The PFPE-DMA showed remarkable robustness and resistance to the e-beam lithography process, enabling nanometer-resolution metal patterning. As a result, this technical advancement allowed for the creation of 32-channel electrode arrays with 500 nm-wide gold interconnects sandwiched by PFPE-DMA dielectric layers (Fig. 5a). BF microscopic imaging verified the precise patterning of the gold interconnects on the PFPE-DMA dielectric layer (Fig. 5b-e). In addition, the high-density stretchable electrode array demonstrated resilience when subjected to stretching, bending, or twisting (Extended Data Fig. 9b).

**Fig. 5.**
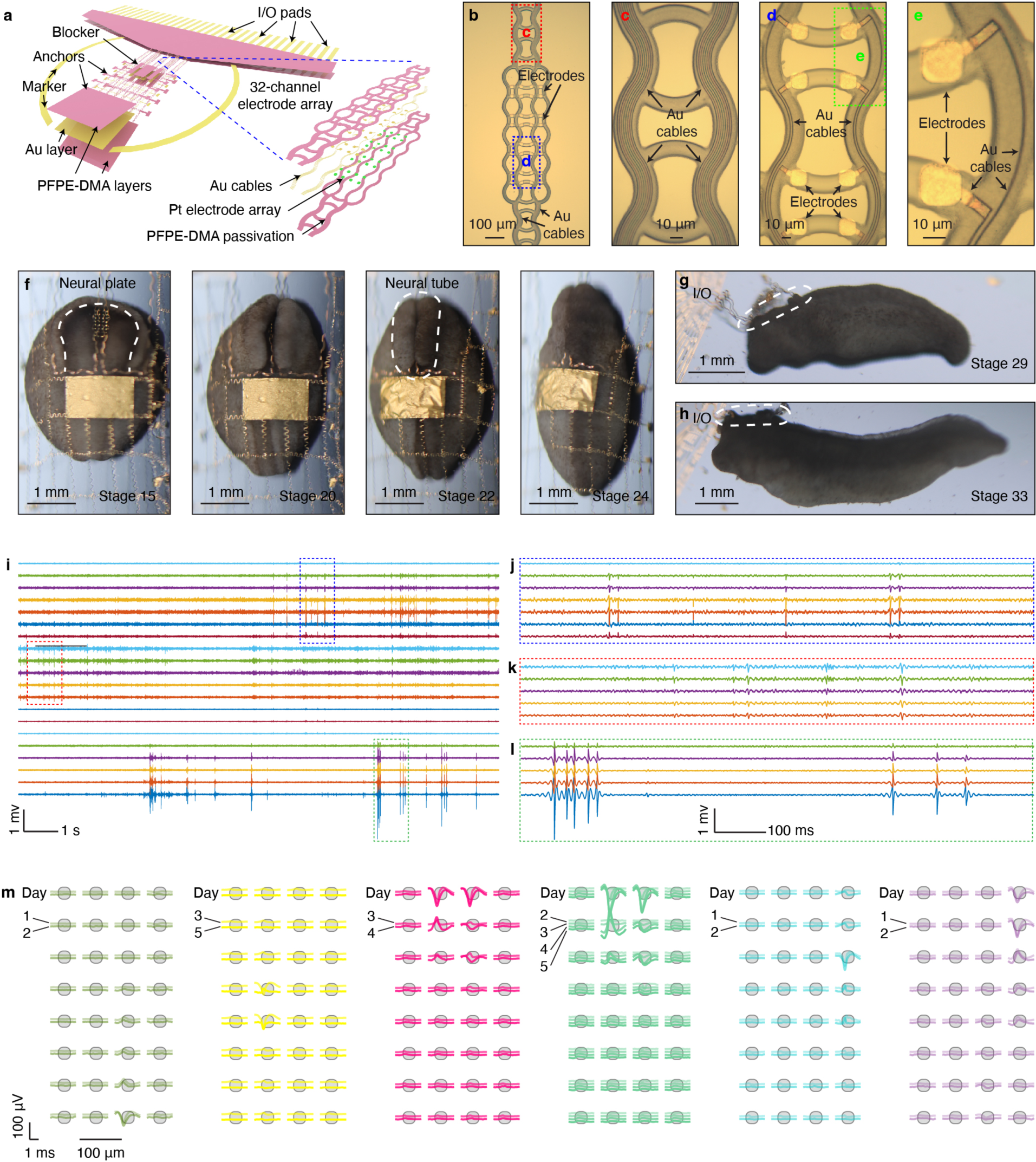
Tracking of single-unit action potential by soft and stretchable high-density mesh electrode array during axolotl embryonic brain development. **a,** Schematic showing the tri-layer structure of a PFPE-DMA-encapsulated stretchable mesh electrode array: PFPE-DMA passivation layers sandwich Au interconnects for a high-density 32-channel electrode array. **b,** BF microscopic image showing the high-density 32-channel electrode array for electrophysiological recording. **c-d,** Zoomed-in view of the red and blue dashed box-highlighted regions in (**b**) showing the stretchable interconnects (**c**) and electrode array (**d**), respectively. **e,** BF microscopic image of the zoomed-in view of the green dashed box-highlighted region in (**d**) showing two individual electrodes. **f,** Time-lapse BF microscopic images of a representative axolotl embryo implanted with stretchable mesh electronics at different development stages, showing the stepwise internalization of the stretchable mesh electronics through the neural plate to neural tube transition (dashed lines). **g, h,** BF images of the axolotl embryo implanted with stretchable mesh electronics at stages 29 (**g**), and 33 (**h**). The dashed circles highlight the stretchable connections between the external interconnects with the implanted electrodes. **i,** Representative filtered voltage traces (300-3,000 Hz bandpass filter) recorded in a cyborg axolotl tadpole. **j-l,** Zoomed-in views of the voltage traces in (**i**) highlighted by blue dashed box (**j**), red dashed box (**k**), green dashed box (**l**). **m,** Representative average single-unit waveforms at each of the recording electrodes over the course of 5-day recording in a cyborg axolotl tadpole.

We used these soft, stretchable high-density mesh electrode arrays to record brain development in the axolotl embryos. Axolotls are an important animal model as their neural system possesses the ability to regenerate after injury. The implantation of these high-density electrode arrays in the brain of axolotl embryos follows the same procedure demonstrated for the *Xenopus* model. Time-lapse BF images (Fig. 5f) show the stepwise internalization of the stretchable mesh electronics by the neural plate during the formation of the neural tube (occurring between stages 15 to 24). Subsequent images captured at stages 29 and 33 (Fig. 5g,h) revealed that as the cyborg axolotl embryo developed into a tadpole, the stretchable meshes were fully embedded within the brain.

We demonstrated that high-density electrode arrays can track single-unit action potentials from the same neurons during axolotl development. Specifically, cyborg axolotl embryos and tadpoles were continuously monitored in culture media with their movements accommodated by the stretchable interconnects of the setup. This allows for continuous recordings up to stage 38. Spike-like signals emerged at stage 25. With interelectrode distances designed to be 50 μm (Supplementary Fig. 5), comparable to the size of a single neuron soma, activity from each neuron could be simultaneously recorded by multiple nearby electrodes^43–45^ (Fig. 5i-l). Sorted spikes and their raster plots across multiple recording days (5-day recording corresponding to stage 27, 28, 29, 31, 33 respectively) showed that similar waveforms could be consistently captured from the same electrode or array of electrodes, validated with interspike interval (ISI) variability (Extended Data Fig. 10a). We overlaid the average spike waveforms recorded by each electrode over multiple days with the layout of the corresponding electrode array. The results showed that spikes originating from one neuron could be consistently recorded by multiple nearby electrodes, with the amplitude varying due to the difference in neuron-electrode distance (Fig. 5m and Extended Data Fig. 10b). Using the spatial arrangement of electrodes and respective mean waveform amplitudes at each electrode, we could estimate the relative position of neurons to the electrode array^43–45^. This resulted in 20 well-isolated individual neurons being continuously recorded (silhouette score^46^ = 0.6047). Using this data, we isolated individual neurons and observed changes in their relative locations to the high-density electrode array during brain development (Extended Data Fig. 9g-i).

## Discussion

We developed novel material, structural, and fabrication methods for the design of tissue-level-soft, stretchable, and embryo-development-compatible electronics. These advances enable a high yield integration (Supplementary Table 3) of the electrode array into the developing brain by leveraging the 2D-to-3D reconfiguration of its nervous system during embryonic development. Immunostaining, fluorescence imaging, gene expression analysis, and behavioral testing yielded no discernable perturbations to brain development or function. The implanted device enabled long-term stable tracking of brain electrophysiology at cellular and millisecond spatiotemporal resolution from the same embryo throughout organogenesis. This work constitutes an entirely novel method for the implantation of soft electronics into a living organism throughout the 3D organ.

We validated our device by continuously recording from the developing embryonic brain in two vertebrate species: frog and axolotl embryos. Frog embryonic recordings showed evolving brain-wide electrical activities during development (Fig. 4o and Extended Data Fig. 8p,q): From stage 20 (Extended Data Fig. 8r-t) to 24, slow-wave synchronized electrical activities propagated across the neural tube from the forebrain to the midbrain. These synchronized signals gradually decoupled as calcium wave-like signals emerged at stage 26, possibly indicating the increasing localization of brain activity. By stage 40, isolated single-unit action potential-like spikes appeared as frog tadpole brain function matured. These results demonstrate how the frog brain gradually develops localized neural activity during development. Our recordings during axolotl embryonic brain development demonstrated the ability to consistently track the electrical activities from individual neurons and their position changes. Additionally, the recording during spinal cord injury-regeneration showed a significant increase in neuron firing rate during regeneration, which was similar to the activity recorded during the early stages of development when neurons first initiated firing (Extended Data Fig. 10c), suggesting that brain activity was involved in spinal cord regeneration^47^.

In the future, this technology holds the potential for scaling up to higher electrode counts. We pushed the limits of e-beam lithography, showing that 128-channel electrode arrays with 300 nm gold interconnect width can be easily fabricated (Extended Data Fig. 9c-f). Further scalability can be explored through the implementation of multiplexing circuits and 3D multilayer packaging techniques. Given the similarity in neural developmental processes across vertebrates, our technique has the potential for application in other vertebrate species. As preliminary tests, we successfully implanted the stretchable mesh electronics in mouse embryos and neonatal rats and recorded the electrophysiological signals in the developing mammalian brain (Supplementary Figs. 6,7, see Methods). Future experiments with rodent embryos could be conducted using rotator type *in vitro* embryonic culture technology^48^, or with *in utero* culture^49^. Incorporation of electrical or optical stimulators into stretchable meshes could enable the continuous modulation of neural signals during development alongside recording. The integration of sensor registration alongside other *in situ* characterization methods could further enable us to understand how the cellular development gives rise to electrical signals. Additionally, we demonstrated the recording of awake animals by adapting the head-fixation strategy using agarose gel that was previously developed for zebrafish calcium imaging (Supplementary Fig. 8, see Methods). The implementation of virtual reality systems^50^ in conjunction with recording in awake, behaving animals could enable the discovery of behavior- and sensory-specific brain activity throughout development.

## Methods

### 1. Chemicals and materials

All chemicals were obtained from Sigma-Aldrich unless otherwise mentioned and used without further purification. PFPE precursors were obtained from Axoft or Solvay. SEBS were obtained from Asahi Kasei. All photoresists and developers were obtained from MicroChem Corporation unless otherwise mentioned and used without further purification. PFPE-DMA precursors were prepared as previously reported methods^33^.

### 2. Fabrication of stretchable mesh electronics

#### SU-8 devices

Wafer cleaning: A 3-inch thermal oxide silicon wafer (2005, University wafer) was rinsed with acetone, isopropyl alcohol (IPA), water, then blown dry and baked at 110 °C for 3 minutes, followed by O_2_ plasma treatment at 100 W, 40 sccm O_2_ for 30 seconds. (2) Ni sacrificial layer: Hexamethyldisilazane (HMDS) was spin-coated at 4,000 rpm for 1 minute. LOR 3A was spin-coated at 4,000 rpm for 1 minute and hard-baked at 180 °C for 5 minutes. S1805 was spin-coated at 4,000 rpm for 1 minute and hard-baked at 115 °C for 1 minute. Then the photoresists were exposed with 40 mJ/cm^2^ ultraviolet (UV) light and developed with CD 26 for 50 seconds, rinsed with deionized (DI) water, and blown dry. After preparing the photoresist pattern, a 100 nm Ni layer was thermally deposited on the wafer (Sharon) and lifted off in Remover PG for 3 hours. (3) Pt layer: 5/40 nm chromium (Cr)/Pt layers were deposited by electron-beam (e-beam) evaporator (Denton) with photoresist and lifted off in Remover PG for 2 hours. (4) Bottom SU-8 passivation layer: SU-8 2000.5 was spin-coated at 4,000 rpm for 1 minute and pre-baked at 60 °C for 1 minute and 95 °C for 1 minute. SU-8 was exposed with 200 mJ/cm^2^ UV light, then post-baked at 60 °C for 1 minute and 95 °C for 1 minute. SU-8 was then developed in an SU-8 developer for 1 minute, rinsed with IPA, and blown dry. Finally, SU-8 was hard-baked at 180 °C for 1 minute. (5) Au interconnects: 5/50/5 nm Cr/Au/Cr layers were deposited on the top of the bottom SU-8 passivation layer by e-beam evaporator (Denton) with S1805 and lifted off in Remover PG for 8 hours. (6) Top SU-8 passivation layer: The fabrication of the top SU-8 layer followed the same procedure as the fabrication of the bottom SU-8 layer.

#### PFPE-DMA devices by photolithography

Wafer cleaning and the preparation of the Ni sacrificial layer and Pt layer followed the same procedure as the fabrication of SU-8 devices. (3) SU-8 spacers: SU-8 2010 was spin-coated on the wafer at 4,000 rpm for 1 minute and pre-baked at 60 °C for 2 minutes and 95 °C for 2 minutes. SU-8 was exposed with 200 mJ/cm^2^ UV light, then post-baked at 60 °C for 2 minutes, 95 °C for 2 minutes. Finally, SU-8 was developed in an SU-8 developer for 2 minutes, rinsed with IPA, and blown dry. (4) Bottom PFPE-DMA passivation layer: The wafer was first cleaned with acetone, IPA, water, and blown dry. Then the PFPE-DMA precursor was spin-coated at 3,000 rpm for 1 minute and pre-baked at 115 ℃ for 2 minutes. The PFPE-DMA was patterned with 80 mJ/cm^2^ UV in a custom nitrogen chamber, post-baked at 115 ℃ for 2 minutes, developed in developer (bis(trifluoromethyl)benzene: 1,1,1,3,3-pentafluorobutane = 1:3) for 1 minute and blown dry. Finally, the PFPE-DMA pattern was hard-baked at 150 ℃ for 50 minutes. (5) Au interconnects: The PFPE-DMA surface was activated with inert gas plasma for 5 minutes. Then the photoresists, HMDS, LOR 3A, and S1805 were patterned on the wafer as described in the preparation of the Ni sacrificial layer. After that, adhesion metal aluminum (Al) was sputtered at 250 W, 40 sccm argon (Ar) for 90 seconds. Au was sputtered at 125 W, 40 sccm Ar for 3 minutes (AJA International). Finally, the metal layers were lifted off in Remover PG overnight. (6) Top PFPE-DMA passivation layer: Fabrication of the top PFPE-DMA layer followed the same procedure as the fabrication of the bottom PFPE-DMA layer.

#### PFPE-DMA devices by e-beam photolithography

The e-beam fabrication process of PFPE-DMA stretchable mesh electronics followed the same protocol as mentioned above, except for the application of e-beam lithography to pattern Au interconnects on the bottom PFPE-DMA layer. Specifically, after the PFPE-DMA surface was activated using inert gas plasma for 5 minutes, the e-beam resist methyl methacrylate (MMA) EL7 was spin-coated at 4,000 rpm for 1 minute and then hard-baked at 150°C for 90 seconds. Next, the e-beam resist 950 polymethyl methacrylate (PMMA) A6 was spin-coated at a rate of 4,000 rpm for 1 minute and then hard-baked at 180°C for 90 seconds. A 10 nm layer of Au was sputtered (AJA International) to assist the e-beam lithography process on the PFPE-DMA. The e-beam resists were patterned at 1000 µC/cm^2^ (Elionix ELS-HS50). The Au layer was then removed using Au etchant, and the resists were developed (methyl isobutyl ketone (MIBK):IPA = 1:3) for 1 minute. Final steps included sputtering adhesion metal Al at 250 W, 40 sccm Ar for 90 seconds, and Au at 125 W, 40 sccm Ar for 3 minutes (AJA International). The process was completed with an overnight lift-off of the metal layers in Remover PG.

### 3. Characterizations

#### Three-point bending test

10 g/mL gelatin (G1890) was dissolved in 60 ℃ DI water. The gelatin solution was then cooled to 25 ℃ to form a thin membrane. PFPE-DMA mesh electronics were laminated on the membrane. Additional gelatin solution was added and cooled to embed the mesh electronics. Gelatin membranes were adhered over a 1-mm-wide gap on 3D-printed plastic substrates. An AFM cantilever (BRUKER, SAA-SPH-1UM) was loaded onto the center of the samples over the gap. The force and displacement of the cantilever were recorded.

#### Elastic modulus measurement

The contact mode of an AFM (JPK Nanowizard AFM) was used to measure the elastic modulus of *Xenopus* embryonic tissue and organoids. Embryonic tissues were tested in 0.1×MMR (a 1 L H_2_O solution containing 5.844 g NaCl (S7653), 0.1492 g KCl (P3911), 0.1204 g MgSO_4_ (M7506), 0.2940 g CaCl_2_ (C1016), 1.192 g 4-(2-hydroxyethyl)-1-piperazineethanesulfonic acid (HEPES) (H3375), 200 mg gentamycin (VWR International, 0304), and 100 mg NaOH (S8045)), and brain and cardiac organoids were tested in 1×phosphate-buffered saline (PBS) (VWR international, 97063-660). Samples were secured using custom 3D-printed parts specifically designed to fit the sample shape and maintain stability during measurements. The AFM cantilever (BRUKER, SAA-SPH-1UM) was loaded to record the force and displacement for the calculation of the elastic modulus.

#### Contact angle and surface free energy

Contact angles for various liquids (water, diiodomethane (158429), and phospholipid (P3817)) were measured through contact angle measurement techniques. Images were analyzed by using Fiji to determine the contact angle. The surface free energy of each substrate was calculated from the contact angles of water and diiodomethane using the Fowkes model^51^.

### *In vitro* biocompatibility test

The PFPE-DMA mesh electronics were heated at 80 ℃ in a 1 M NaOH solution for 2 hours to obtain its potential degradation products. After being washed by DI water 10 times to remove the NaOH solution, PFPE-DMA devices or degradation products was mixed with the culture medium at a concentration of 1% v/v and co-cultured with wild-type rat cortical neurons for 10 days. The live cell ratio of the neurons was compared to the control using a cell viability and cytotoxicity assay (CELL BIOLABS, CBA-240).

### 4. Animal experiments

Frog embryos were obtained from the National Xenopus Resource. Axolotl embryos were obtained from the University of Kentucky (AGSC_100E). Mouse embryos were collected from pregnant C57BL/6 mice (Charles River Laboratories INC) based on previous reports^52,53^. Neonatal rats were bred from pregnant CD rats (Charles River Laboratories INC) following previously reported protocols^54^. All experimental procedures were approved by the Institutional Animal Care and Use Committee (IACUC) of Harvard under animal protocols # 19-01-344-1 and # 19-03-348-1.

#### Device assembly and treatment for implantation

Wafer was cut using a dicing saw, protected by photoresist S1813 during the cutting process, and the photoresist was removed afterwards. (2) A flexible flat cable (FFC) (Molex) was bonded to the I/O pads using a flip-chip bonder (Finetech Fineplacer). (3) A microscope slide (VWR International, 48300-026) was used as a stable base, adhered to the wafer piece with low toxicity silicone adhesive (World Precision Instruments, KWIK-SIL). A 50 mL centrifuge tube (VWR International, 525-0610) was cut to serve as the culture chamber and attached to the wafer piece with the same silicone adhesive. (4) The device was released from the substrate by removing the Ni sacrificial layer in Ni etchant (TFB, Transene Company), which was subsequently washed out using DI water for 10 times. The device was then incubated with 1 mL 0.01% poly-D-lysine hydrobromide solution (P4832) overnight. (5) The device was washed with DI water three times and then incubated with 1 mL 10 mg/mL culture media diluted Matrigel (Corning, 08-774-552). Culture media for frog embryos was 0.1×MMR; for axolotl embryos was 1× Steinberg’s solution (1 L H_2_O solution contains 0.34 g NaCl, 0.005 g KCl, 0.008 g Ca(NO_3_)_2_·4H_2_O (C2786), 0.01025 g MgSO_4_ (M2643), 0.056 g Tris-HCl (10812846001), 0.001 g phenol red (P3532)); for mouse embryos was rat serum prepared as in previous reports^52,53^.

#### Implantation of frog and axolotl embryos

De-jellying: Frog embryos need de-jellying before implantation. Embryos were placed in a de-jelly solution (60 mL 1×MMR containing 1.2 g L-Cysteine (168149) and 0.1 g NaOH) for 5 minutes and then washed with 1×MMR five times. (2) Implantation: A stage 15 embryo was placed inside the culture chamber under a stereoscope. Next, the vitelline membrane of the embryo was peeled off using #5 tweezers (Fine Science Tools, 11252-40) to expose the neural plate. The embryo was then slid under the stretchable device using tweezers, ensuring that the implanted ribbons accurately overlapped with the neural plate. During this process, one tweezer held the device while another guided the embryo into position underneath.

#### Implantation of mouse embryos

Prior to implantation, the culture well plate was transferred from the incubator to a 37°C heating pad. A pre-sterilized wafer piece carrying the anchored device was immersed in the media. Then the embryo was placed on the wafer, lying on its side with the neural plate facing the device. Using tweezers, the device was carefully elevated. Subsequently, the embryo was gently pushed toward the device to implant the device into the neuropore. Finally, the anchor was cut, and the wafer piece was gently removed. The entire process needed to be completed within 10 minutes, and the embryos were promptly returned to culture immediately after implantation. The culture protocol of mouse embryos was adapted from previous reports^52,53^.

#### Stereotaxis implantation of neonatal rats

Rat pups were kept on a regular 12-hour light-dark cycle and housed with their mother before surgery. Pups underwent implantation at P5-7. They were placed on a customized platform for head fixation and were maintained under anesthesia with 0.5% isoflurane during surgery. For implantation, a circular piece of skin on the skull was cut away with surgical scissors. Then, a small cranial window was opened, and the probe was stereotactically advanced until all electrodes were inserted below the pial surface. A 100 µm-thick stainless-steel wire (A-M SYSTMES, 793100) was partially inserted to the window as the ground. The device’s I/O and the ground were carefully sealed and fixed with dental cement. After implantation, the rat pups were placed back with their mother. Postoperative care was provided. Carprofen should be administered to rat pups for the following 4 days, both in the morning and in the afternoon. Activity, incision, and pain were monitored and recorded daily after implantation.

#### Long-term rearing of cyborg frog tadpoles

Control and experimental groups were split evenly between two static 5-gallon tanks with comparable conditions. They were reared in 0.1xMMR solution with partial water changes completed 2-3 times weekly. During water changes, evaporative loss was compensated for, and the pH was buffered with alkaline buffer (Seachem Laboratories, Inc.), as needed. Biological filtration was developed prior to animal introduction using sponge filters. Animals were fed Sera micron (Sera North America Inc.) and Fry Starter (NorthFin Fish Food). Tanks were spot siphoned between water changes. Animal health was checked daily, and water quality was analyzed at least once a week to ensure it was within healthy parameters.

#### Behavior tests of frog tadpoles

Tadpoles for behavior tests were cultured at room temperature (RT) on a white base under a 12-hour day-night cycle. Behavior tests were conducted between 1 and 4 pm. For each trial, a tadpole was placed at the center of a clear-bottomed round tank (diameter of 12 cm) filled to 5 cm with 0.1×MMR. The tank was positioned on a horizontal screen and covered in a dark box. The luminance of the screen was set to 50 cd/m^2^ when displaying the white color. During the test, the tadpole was stimulated by the appropriate pattern shown on the screen. In the color preference test, the screen alternated between displaying half white and half black for 40 seconds each. In the visual avoidance test, a black dot was moved towards the tadpole on the screen. In the edge preference test, the tadpole swam freely on the white screen. The tadpole’s response was video recorded by a camera positioned on top of the dark box. Any disturbances, including vibrations, light, and sounds, needed to be avoided.

#### Whole-mount staining of cyborg frog tadpoles

Fixation and bleaching: Tadpoles were fixed with 4% paraformaldehyde (PFA) (Thermo Fisher SCIENTIFIC, J19943-K2) overnight at 4 ℃ and then transferred to a glass dish containing a bleaching solution: (1.5 mL 30% H_2_O_2_ (H1009), 2 mL formamide (47670), 1 mL 20×SSC buffer to 35 mL DI). The dish was placed on a nutator over aluminum foil reflective backing and under fluorescent light. To remove bubbles caused by bleach, embryos were dehydrated in methanol for 5 minutes. Subsequently, the embryos were rehydrated over the course of 10 minutes in stages of 80% methanol / 20% DI water; 50% methanol / 50% PBS; 20% methanol / 80% PBS. (2) Staining: Embryos were washed with 0.1% PBST (50 mL PBS contains 50 μL Triton X-100 (X100-1L)) twice, 30 minutes per wash. Then, they were incubated in diluted CAS-Block (13.5 mL PBS containing 1.5 mL CAS-Block (Thermo Fisher SCIENTIFIC, 008120)) for 1 hour at RT. After that, tadpoles were stained in primary antibody solution (1 mL CAS-Block containing 10 μL anti-acetylated tubulin (T7451) and 10 μL anti-HuC/D (Abcam ab184267)) for 2 days at 4 ℃. Then the embryos were washed with PBST for 30 minutes at RT and then blocked in PBST-CAS for 30 minutes at RT. Subsequently, the embryos were incubated in secondary antibody solution (1 mL CAS-Block containing 2 μL Alexa Fluor 488 (Invitrogen, A-11006), 2 μL Alexa Fluor 594 (Invitrogen, A-11012), 20 μL Alexa Fluor 647 Phalloidin (Thermo Fisher SCIENTIFIC, A22287) and 1 μL DAPI (D9542)) for 2 days at 4 ℃. Finally, the embryos were washed with PBST for 1 hour at RT and then washed with PBS overnight at 4 ℃. (3) Imaging: Stained tadpoles were imaged in a homemade chamber. A 2 mm layer of vacuum grease (Z273554) was applied at the edge of a microscope slide to form the walls of a chamber. A tadpole was placed in the middle of the microscope slide and embedded in the mounting medium (Vector, H-1900). A cover glass was placed on the top of the vacuum grease to seal the chamber. The tadpole was positioned upside down on the confocal microscope (Leica, dMi8) and imaged with 5 μm stacks. All the images were stitched and processed with Fiji.

#### Cryosection staining of cyborg frog tadpoles

Fixation: Tadpoles were fixed with 4% PFA overnight at 4 ℃ and then incubated in 0.1% PBST overnight at 4 ℃, followed by incubation in 0.1% PBST containing 15% gelatin (G1890)/15% sucrose (S7905) overnight at 40 ℃. Subsequently, the tadpoles were frozen in the gelatin/sucrose solution at -80 ℃ overnight for cryosection. (2) Staining: Slides were placed in a wet box and incubated at 40 ℃ to remove residual gelatin/sucrose solution. Then, the slides were washed with 0.1% PBST for 15 minutes at RT and incubated in blocking buffer (20 mL 0.1% PBST containing 0.2 mL donkey serum (Jacksonimmuno, 017-000-121) and 0.8 g bovine serum albumin (Thermo Fisher SCIENTIFIC, BP1600-100)) for 1 hour at RT. Subsequently, slides were incubated in primary antibody solution overnight at 4 ℃. Slides were then washed with 0.1% PBST three times and incubated in secondary antibody solution overnight at 4 ℃. Finally, the slides were washed with 0.1% PBST three times and sealed. Primary antibody solution for Sox2, Myt1, and acetylated tubulin staining: 2 mL blocking buffer containing 20 μL anti-Sox2 (Invitrogen, 14-9811-82), 20 μL anti-Myt1 (Abcam, ab251682) and 20 μL anti-acetylated tubulin (T7451). Secondary antibody solution for anti-Sox2, anti-Myt1, and anti-acetylated tubulin staining: 4 mL blocking buffer containing 8 μL Alexa Fluor 488 (Invitrogen, A-11006), 8 μL Alexa Fluor 594 (Invitrogen, A-11012), 8 μL Alexa Fluor 647 (Invitrogen, A-32787) and 4 μL DAPI (D9542). Primary antibody solution for BrdU staining: 2 mL blocking buffer containing 20 μL BrdU monoclonal antibody (Invitrogen, B35128). Secondary antibody solution for BrdU staining: 4 mL blocking buffer containing 8 μL Alexa Fluor 647 and 4 μL DAPI. Primary antibody solution for *Aldh1l1* and *vimentin* staining: 2 mL blocking buffer containing 20 μL anti-Aldh1l1 antibody (Abcam, AB56777) and 20 μL anti-vimentin (Abcam, AB16700). Secondary antibody solution for *Aldh1l1* and *vimentin* staining: 4 mL blocking buffer containing 8 μL Alexa Fluor 594, 8 μL Alexa Fluor 647 and 4 μL DAPI.

#### Cryosection staining of cyborg mouse embryos

Fixation: Embryos were fixed with 4% PFA at 4 ℃ overnight, then soaked in 10% sucrose solution at 4 ℃ overnight; followed by 20% sucrose solution at 4 ℃ overnight. The soaked embryos were positioned in a cube chamber with O. C. T. compound (Tissue-Tek 4583) and frozen at -80 ℃ overnight for cryosection. (2) Staining: Before staining, slides were placed in a wet box and incubated at 40 ℃ to remove residual O. C. T. compound. Then the slides were washed with 0.1% PBST for 15 minutes at RT and stained with 4 mL 0.1% PBST containing 4 μL DAPI overnight at 4 ℃. Finally, slides were washed with 0.1% PBST three times and sealed.

#### Confocal imaging

Stained tissue slides were imaged under all samples were imaged with a Leica TCS SP8 confocal microscope using the Leica Application Suite X software platform 3.5.5 (https://www.leica-microsystems.com/products/microscope-software/p/leica-las-x-ls/downloads/). All the images were processed with Fiji.

#### Electrophysiology recording

All recordings were taken with a Blackrock CerePlex Direct recording system or an Intan RHD recording system. For data acquisition, a Blackrock Microsystems CerePlex μ headstage or an Intan RHD recording headstage was connected to the flat flexible cable through a laboratory-made printed circuit board. The recording setup was placed on an optic table and covered by a Faraday cage. For awake and behaving animal recording, cyborg tadpoles were head-fixed by a previously reported agarose fixation method^55,56^. Specifically, a layer of agarose was cured on the agar scaffold of a head-fix setup using an agar mold (**Supplementary Fig. 8a, step 1**). Then, the setup was placed on top of the culture chamber (**Supplementary Fig. 8a, step 2**). Next, the tadpole was fixed using a small amount of low melting point agarose, with the scaffolds serving as anchor points (**Supplementary Fig. 8a, step 3**). The agarose was carefully trimmed to avoid covering the tadpole’s mouth or tail, allowing it to breathe and move its tail once it recovered from the anesthesia. After fixation, the lid was placed on top of the setup to reduce media evaporation. Half of the media was changed every 12 hours to increase the survival rate of the fixed tadpole.

## 5. Data analysis

### Analysis of fluorescence images

Cell counting in Sox2 and Myt1 images: DAPI was used to identify the cell nuclei for accurate cell counting. Using Sox2 as an example, Sox2 image was first binarized to identify the region of neuron stem cells (**Supplementary Fig. 4a, step 1**). The binary Sox2 image was then overlaid with the DAPI image (**Supplementary Fig. 4a, step 2**), and DAPI-labeled cell nuclei in the Sox2 region were counted as the cell number of neuron stem cells (**Supplementary Fig. 4a, step 3**). (2) Fluorescent intensity of BrdU images: The DAPI image was binarized to identify the tissue region (**Supplementary Fig. 4b, step 1**). The binary DAPI image was then overlaid with the BrdU image (**Supplementary** Fig. 4b**, step 2**), and the fluorescent intensity of BrdU within the binary DAPI region was calculated (**Supplementary Fig. 4b, step 3**). (3) Fluorescent intensity of *Aldh1l1* and *vimentin* images was analyzed using Fiji.

### Analysis of behavior tests

The behavior videos were processed by MATLAB to analyze the trajectory of the tadpoles. (1) Color preference test: The tadpole’s distance to white/black boundary was calculated and reported as “tadpole to middle line” (**Extended Data Fig. 6j**). The percentage of time that the tadpole stayed in the white side of the tank was calculated and reported as the “time ratio in white side”, and statistical results of all tested tadpoles were documented in **Fig. 3o**. (2) Visual avoidance: The tadpole’s distance to the black dot was calculated and reported as “tadpole to black dot” (**Extended Data Fig. 6k**). The percentage of successful escapes from the black dot was calculated and reported as the “avoidance ratio”, and statistical results of all tested tadpoles were documented in **Fig. 3p**. (3) Edge preference: The tadpole’s distance to the tank center was calculated and reported as “tadpole to center” (**Extended Data Fig. 6l**). The percentage of time that the tadpole stayed in the tank edge (the area within 4 cm of the tank wall) was calculated and reported as the “time ratio at the edge”, and the statistical results of all tested tadpoles were documented in **Fig. 3q**.

### Analysis of electrophysiological signals

The electrophysiological recording data were analyzed offline. (1) Filtering: Raw data were filtered using a lowpass filter in the <100 Hz frequency range for oscillation signals, a lowpass filter in the <300 Hz frequency range for calcium wave-like signals, a bandpass filter 300–3,000 Hz frequency range for spike-like signals. (2) Correlation analysis: for signals from stages 24, 26 and local field potential signals in stage 40, one group of four synchronized peaks from each channel were extracted, and Pearson correlations were calculated between each pair of the synchronized peaks. Pearson correlations from all groups of synchronized peaks were then pooled for each stage. Apart from local field potentials, action potentials (spikes) were also extracted for stage 40 signals. Signals chunks from all channels were then extracted according to the time of spikes before the calculation of their pairwise Pearson correlations. Finally, the pooled Pearson correlations for each stage were compared. (3) Synchrony analysis: For signals from stages 24, 26 and local field potential signals in stage 40, the time differences between synchronized peaks in each pair of the channels were collected, generating six datasets for each stage. The standard deviation of each dataset indicated synchronization between two channels. The six standard deviations for each stage were then plotted and compared. (3) Spike sorting: Spike sorting of cyborg frog embryos/tadpoles was performed in MATLAB using WaveClus^57^ and MountainSort^58^. Spike sorting of cyborg axolotl embryos/tadpoles and neonatal rats was performed using a custom pipeline based on SpikeInterface^59^. The spiking time of each single unit were used to compute the ISI histogram.

## 6. Simulations

### FEA analysis

All the FEA were performed using ABAQUS 2022/Standard. (1) Devices: PFPE-DMA and SU-8 ribbons were discretized by S4R element. The material behavior of the ribbons was captured using a linear elastic material model with *E_SUB_* = 6 GPa, *v_SUB_* = 0.4 and *E*_$%$&_ = 500 kPa, *v_PFPE_* = 0.4. The response of the ribbons was simulated by *STATIC module. (2) Embryos: The tissue was discretized by C3D4H element, with finer mesh size toward the neural plate. The material behavior of the embryo tissue was characterized by β_*plate*_ = 150 Pa, β_*ectoderm*_ = 3 kPa and β_*inside tissue*_ = 1.5 Pa. The response of the tissue was captured by an incompressible neo-Hookean material model^60^. (3) Development simulation: A growth model^61^ was used to mimic the embryo development. Volume-proportional damping was added using the option STABILIZE in the ABAQUS STATIC module (dissipated energy fraction equal to 5 × 10^67^ and the maximum ratio of stabilization to strain energy equal to 0.05). The development was driven by a thermal expansion of the embryo tissue, relating to the thermal strain ε^*th*^ through ε^*th*^ = α(θ − θ^*I*^), where α was the thermal expansion coefficient of the material, θ was the current temperature and θ^*I*^ was the initial temperature. In our simulations, ε_*plate*_ = 0.4 [1/K], ε_*inside tissue*_ = 0.6 [1/K], ε_*ectoderm*_ = 1.0 [1/K], the temperature *θ* was gradually increased until the neural plate formed a deep fold and encapsulated the device.

## Acknowledgments

J.L. acknowledges the support from the Startup fund from the School of Engineering and Applied Sciences, Harvard University; National Institutes of Health Award NIH/NIMH 1RF1MH123948; Aramont Fund for Emerging Science Research; and the William F. Milton Fund. H.S. acknowledges the support from Aramont Fund for Emerging Science Research. The authors thank Daniel J. Needleman and Jessica L. Whited for their helpful discussions.

## Author contributions

J.L. and H.S. conceived the idea. R.L., H.S., P.L., W.W., and A. L. fabricated and characterized electronics. H.S. performed implantation and behavior tests. H.S. and Q.L. applied immunofluorescence. H.S., Z.L. and J.B. did qPCR. H.S. and R.L. performed electrical recording. L.J. and Z.W. did the mechanical simulation. R.J. housed embryos to frogs. H.S., H.Z., Z.L., Y.H., X.T., T.S.B., D.S., and S.Z. analyzed data. All authors prepared figures and wrote the manuscript. J.L. supervised the study.

## Additional information

Correspondence and requests for materials should be addressed to Jia Liu.

## Extended Data Figures and Figure Legends

**Extended Data Fig. 1.**
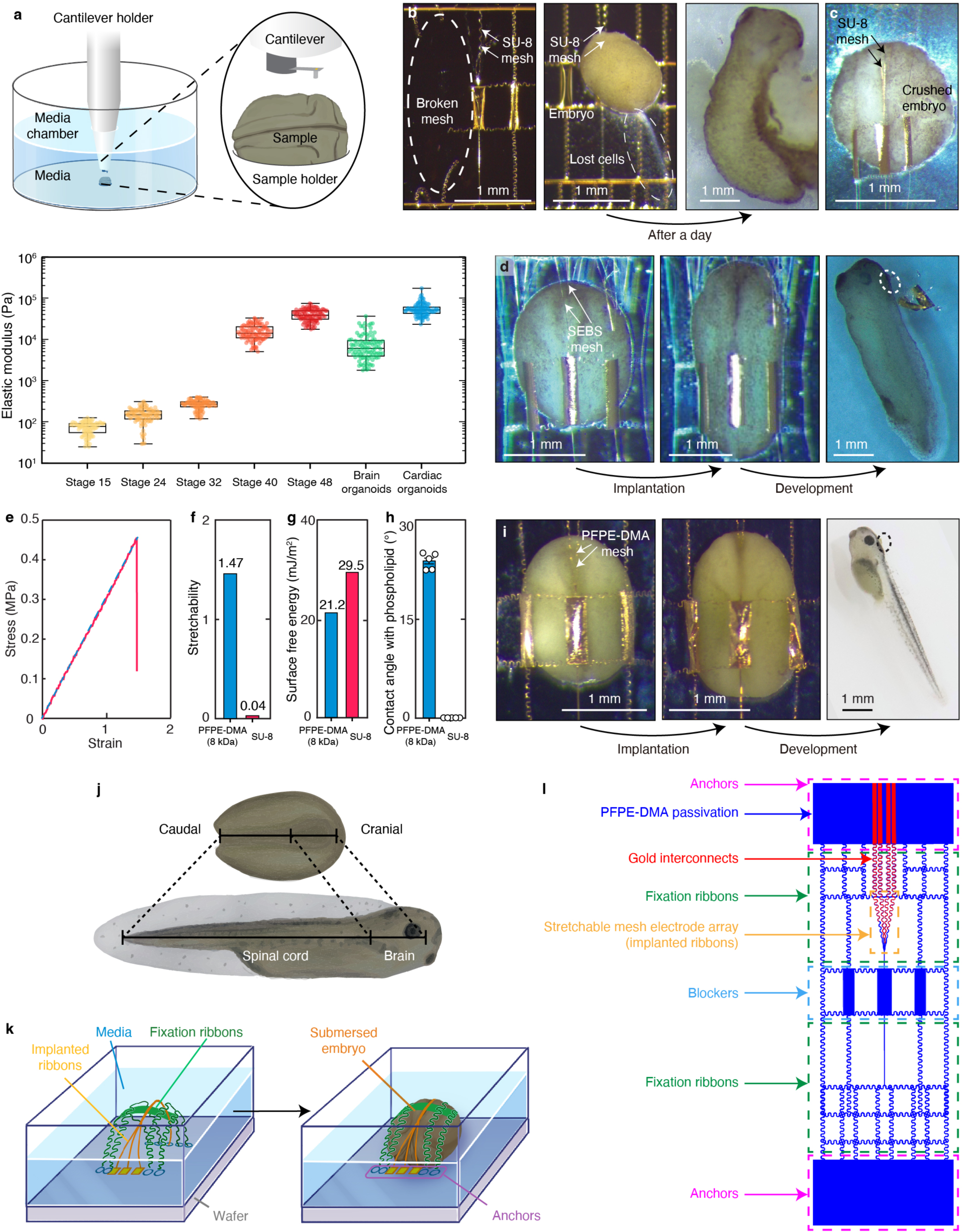
Test of implantation methods. **a,** (Top) schematics showing an atomic force microscopy (AFM) setup for tissue elastic modulus measurement. (Bottom) elastic modulus of stage 15, 24, 32, 40, 48 Xenopus embryos, brain organoids, and cardiac organoids. Box plots indicate minimum, lower quartile, median, upper quartile, and maximum. Each dot represents a contact measurement. **b,** Photographs showing broken SU-8 mesh post-implantation (left) and the embryo before (middle) and after (right) mesh implantation, depicting damage to the embryo. **c,** BF microscopic images showing an embryo crushed by SU-8 meshes. **d,** BF microscopic images showing an embryo successfully implanted with a SEBS mesh. The dashed line circle highlights the portion of the mesh which remains exterior to the tadpole brain. **e,** Stress-strain curve of PFPE-DMA film with 8 kDa molecular weight, the blue dashed line indicates a linear relationship. **f,** Stretchability of 8 kDa PFPE-DMA and SU-8 films. **g,** Surface free energy of 8 kDa PFPE-DMA film and SU-8 films. **h,** Contact angle of phospholipid (cell membrane analog) on 8 kDa PFPE-DMA film, and on SU-8 film. **i,** Photographic images showing an embryo successfully implanted with a PFPE-DMA mesh. The dashed line circle highlights the portion of the mesh which remains exterior to the tadpole brain. **j,** Schematics showing elongation of the neural tube during the embryo development of *Xenopus laevis*. The caudal region of the neural tube elongates to 3 times its initial length and forms the spinal cord while the cranial region elongates only 1.3 times its initial length and forms the brain. **k,** Schematics showing how anchors fix the stretchable mesh electronics to the substrate, keeping the neural plate properly positioned during neurulation for device internalization, and keeping the stretchable mesh electrode array attached to the neural plate. The device’s initial dimensions and stretchability enable the stage 15 embryo to be slid under the device for implantation. **l,** The design of the stretchable mesh electronics showing the architecture of the stretchable mesh electrode array, stretchable serpentine interconnects, anchors, stretchable ribbons, and blockers. The blocker prevents the mesh electrodes from implanting into the caudal region of the neural plate.

**Extended Data Fig. 2.**
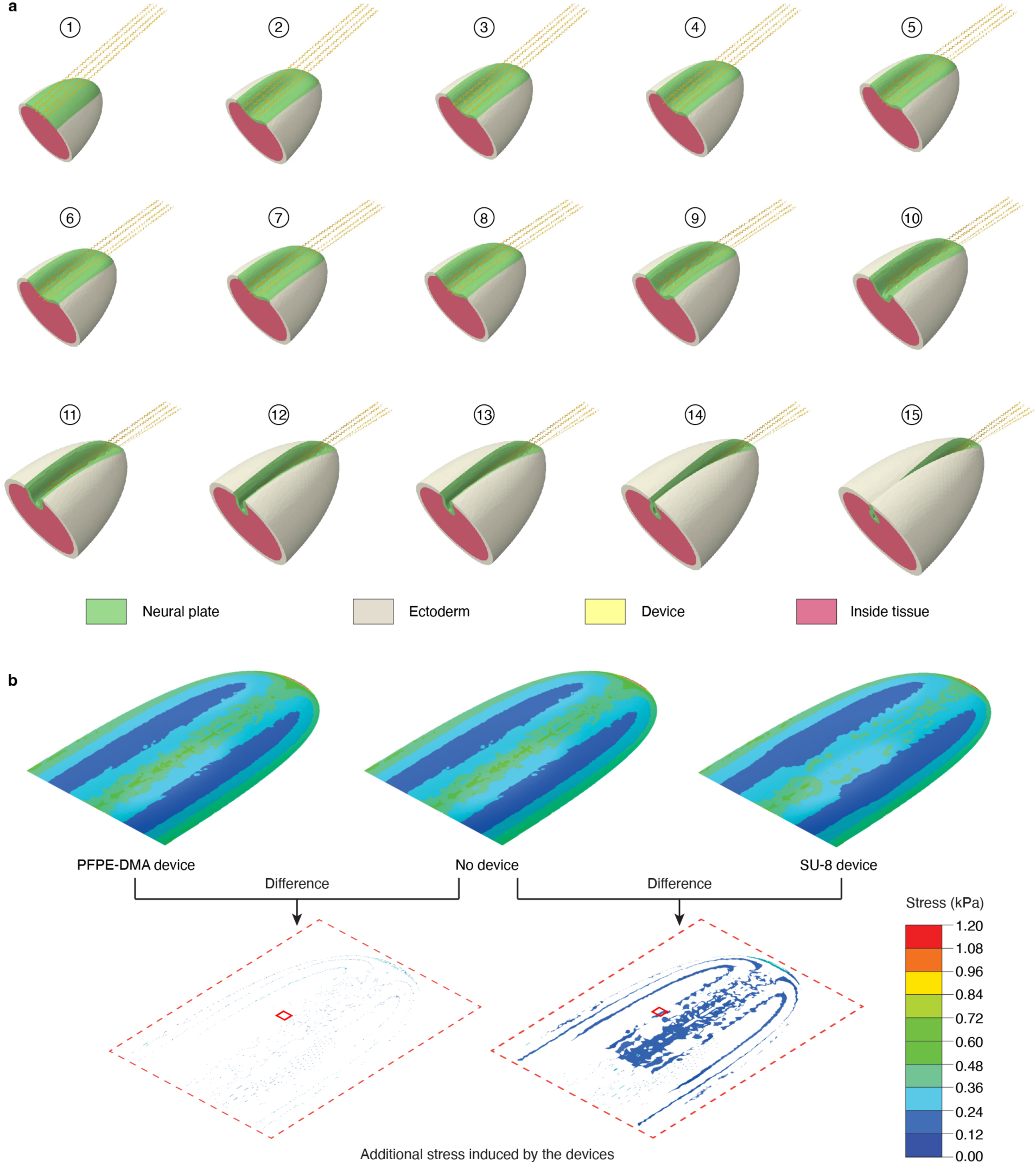
Mechanical simulation of stretchable mesh for brain implantation via embryo development. **a,** Snapshots of mechanical simulation of mesh-neural plate interaction (Fig. 1j, k), labeled with sequenced numbers. **b,** Snapshots of mechanical simulation procedure showing the stress distribution in the neural plates with and without stretchable mesh implanted. An embryo simulation without mesh implantation was used as a reference to calculate the additional stresses introduced by PFPE-DMA and SU-8 meshes. The red boxes highlight regions where the maximum stress was shown in (Fig. 1k).

**Extended Data Fig. 3.**
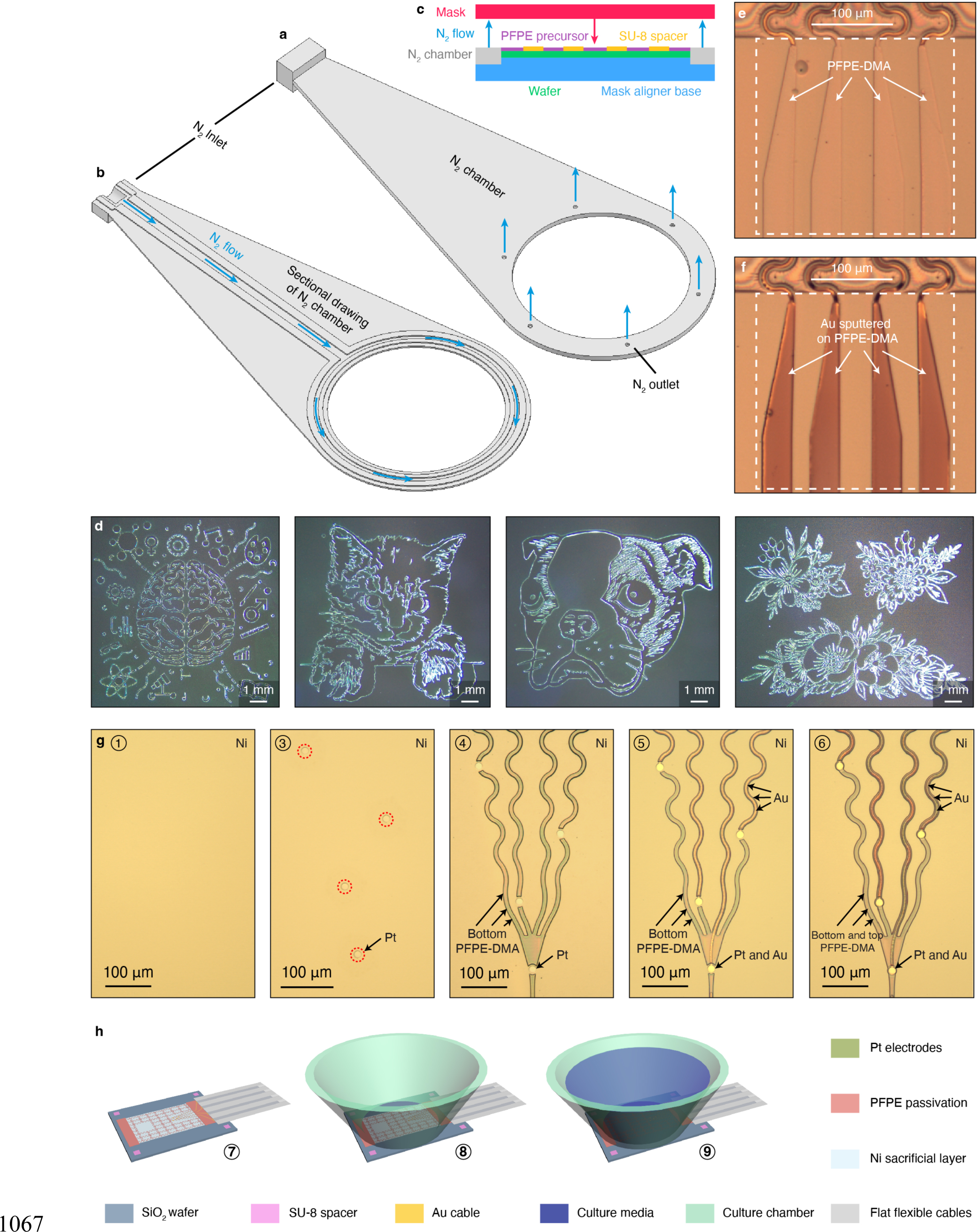
Fabrication of PFPE-DMA-encapsulated stretchable mesh electronics. **a, b,** Schematics showing the overlook (**a**) and section view (**b**) of the nitrogen chamber designed for use with the mask aligner in PFPE-DMA photopatterning. **c**, Schematic showing how the nitrogen chamber is used with mask aligner. **d,** Microscopic BF images showing representative high-resolution PFPE-DMA photolithography patterns made with the nitrogen chamber. **e, f,** Microscopic BF images showing the improved adhesion between Au interconnects and PFPE-DMA after inert gas plasma treatment. Dashed boxes highlight the sputtered regions on the PFPE-DMA layers. Without inert gas plasma treatment, Au interconnects peel off from the PFPE-DMA film after sputtering (**e**). With inert gas plasma treatment before sputtering, Au interconnects strongly bond to the PFPE-DMA film (**f**). **g,** Microscopic BF images showing the stretchable mesh electrode array region of PFPE-DMA device in fabrication steps corresponding to (Fig. 2c). Step 1 shows a homogeneous Ni layer. Step 2 is not included because the electrode array region does not have an SU-8 spacer. Step 3 shows Pt electrodes on the Ni layer. Electrodes are highlighted by red dashed circles. Steps 4-6 show sequential patterning of bottom PFPE-DMA, Au interconnects, and top PFPE-DMA layers. **h,** Schematics showing the post-fabrication steps of PFPE-DMA-encapsulated stretchable mesh electronics following (Fig. 2c). After fabrication, the device is soldered with a flexible flat cable (step 7) and bonded with a culture chamber (step 8). Then, the Ni layer is etched to release the device. Pt-black is electro-polymerized on electrodes to reduce electrode impedance. The device is washed with 0.1 × MMR and finally soaked in culture media (step 9).

**Extended Data Fig. 4.**
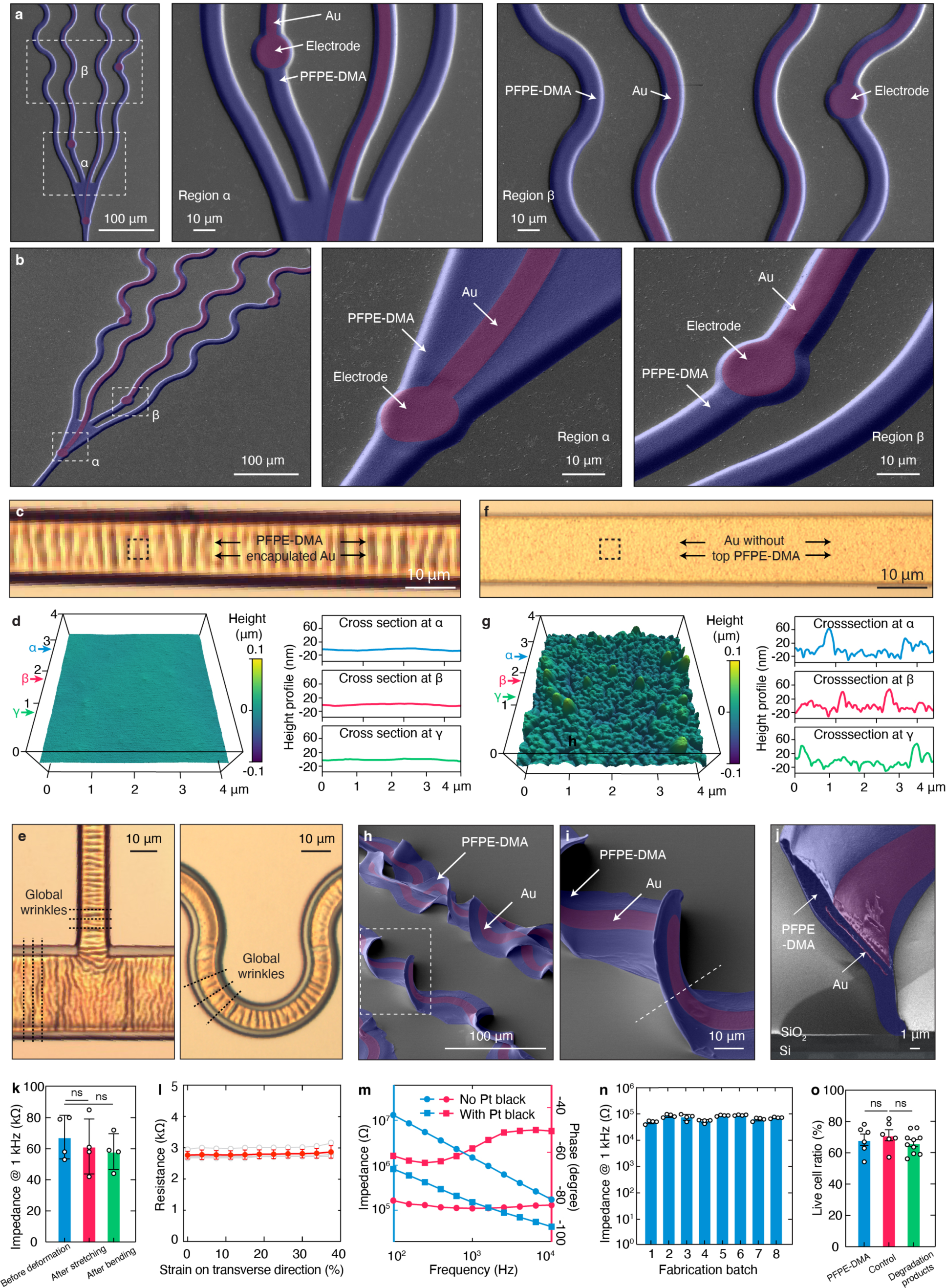
Characterization of PFPE-DMA-encapsulated stretchable mesh electronics. **a, b,** SEM images showing top views (**a**) and perspective views (**b**) of the stretchable mesh electrode array portion of the PFPE-DMA device. Each layer is pseudo-colored and labeled. **c,** BF image of a PFPE-DMA-encapsulated Au ribbon. **d,** (Left) atomic force microscopy (AFM) topography image of black dashed box-highlighted region in (**c**). (Right) height profiles of horizontal cross-sections highlighted in the left figure. **e,** BF images showing wrinkles of (left) straight and (right) serpentine PFPE-DMA-encapsulated interconnects. **f,** BF image of an Au interconnect without top PFPE-DMA passivation. **g,** (Left) AFM topography image of black dashed box-highlighted region in (**f**). (Right) height profiles of horizontal cross-sections highlighted in the left figure. **h,** SEM image showing perspective views of the PFPE-DMA stretchable mesh electronics after stretching and bending. **i,** SEM image of the dashed box-highlighted region in (**h**). **j,** SEM image showing cross-sections of PFPE-DMA-encapsulated Au interconnects, along the dashed line in (**i**). **h-j,** Each layer is pseudo-colored and labeled. **k,** Electrode impedance at 1 kHz in 37 °C PBS of PFPE-DMA mesh electronics before and after stretching and bending. Bar plots indicate mean ± s.e.m., each dot represents a single trial. two-tailed unpaired t-test, *n* = 4, ns, not significant. **l,** Resistance as a function of strain during the transverse stretch test of PFPE-DMA-encapsulated electronics. Red dots and line plots indicate mean ± s.d., and each gray dot and line plot represents one sample. **m,** Electrochemical impedance spectroscopy of electrodes in stretchable mesh electronics with and without Pt-black coating. **n,** Electrode impedance at 1 kHz in 37 °C PBS of PFPE-DMA mesh electronics fabricated in different batches. **o,** Live cell ratio of wild-type rat cortical neurons after 10 days *in vitro* culture with PFPE-DMA mesh electronics, control, and with degradation products of PFPE-DMA. Bar plots indicate mean ± s.e.m., each dot represents a single trial. two-tailed unpaired t-test, *n* = 6, ns, not significant.

**Extended Data Fig. 5.**
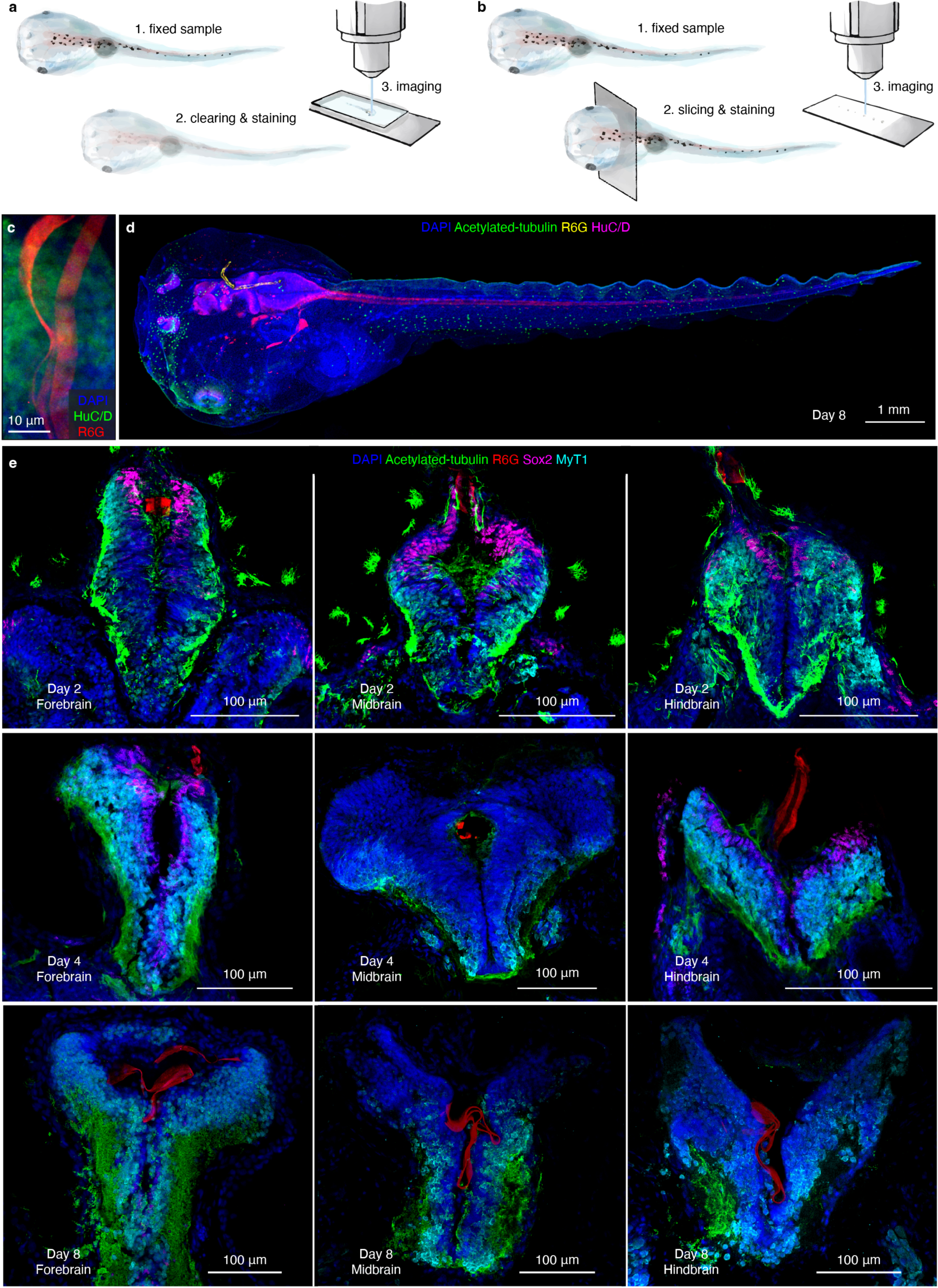
Staining methods and extended immunofluorescence images. **a, b,** Schematics showing the protocols for tissue clearing and whole-mount staining (**a**) and cryosection staining (**b**) to characterize brain tissue implanted with stretchable mesh electronics. **c,** Whole-mount-stained 3D reconstructed confocal fluorescence image of implanted mesh electronics showing that the mesh is embedded in the neural tissue. **d,** 3D reconstructed confocal fluorescence images of a whole-mount-stained cyborg tadpole whose device was implanted in the middle of neurulation. **e**, Confocal fluorescence images showing coronal sections of the fore-, mid-, and hindbrain of cyborg tadpoles fixed at 2-, 4- and 8-days post fertilization. In all images, DAPI labels cell nuclei, acetylated-tubulin labels basal bodies, R6G labels the device, and SRY-box transcription factor 2 (Sox2) labels neural stem cells. In fluorescence images of whole-mount staining samples, HuC/D labels neurons. In fluorescence image of cryosection staining sample, Myt1 labels neurons.

**Extended Data Fig. 6.**
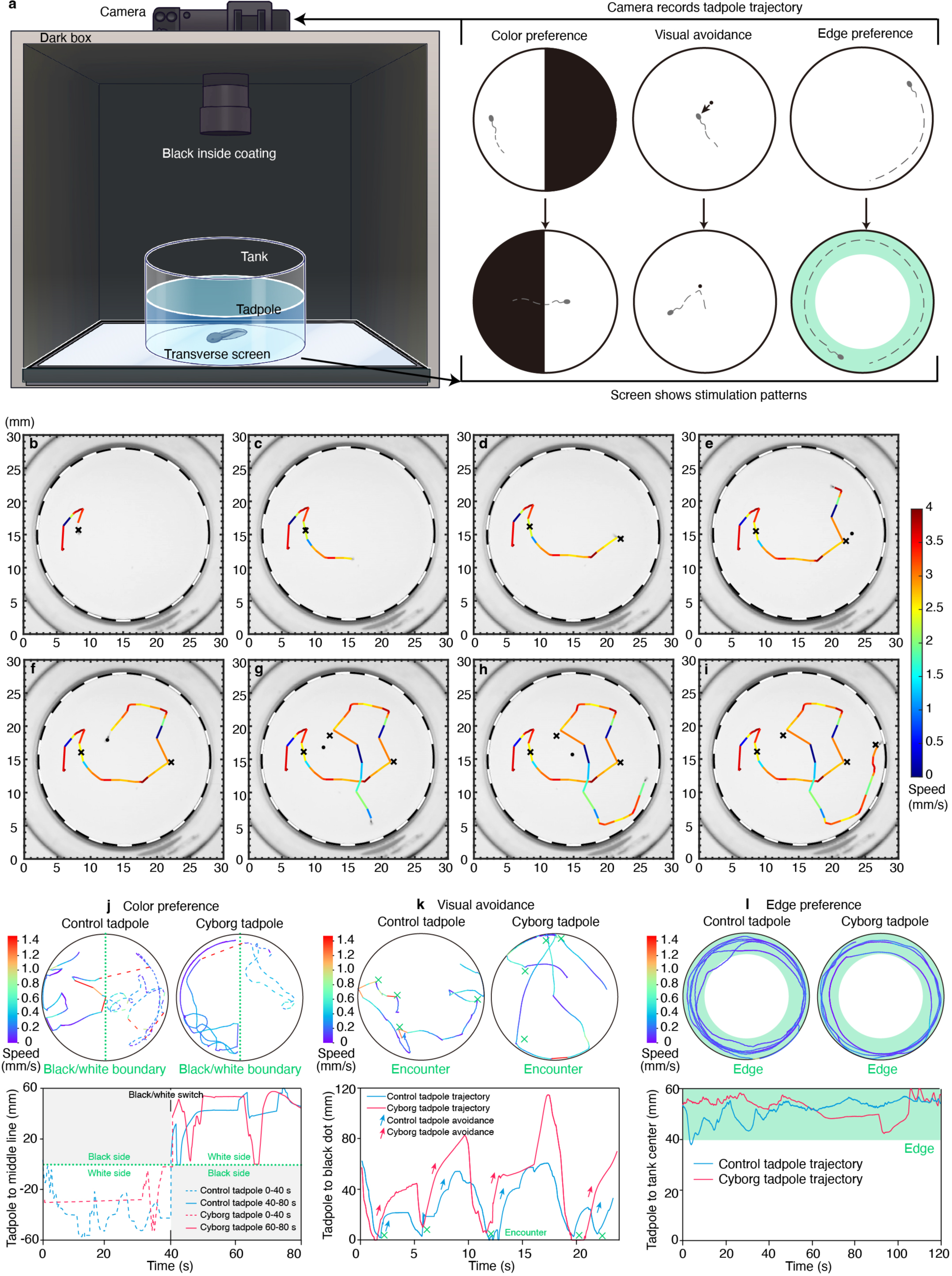
Experimental setup, trajectory analysis, and examples of behavior tests. **a,** Schematics showing the setup for behavioral testing. In each test, a tadpole is placed in a clear tank, sitting on an upward-facing screen. The screen is programmed to display the appropriate stimulation pattern for the color preference, visual avoidance, and edge preference tests. In the color preference test, the screen alternated between displaying half white and half black for 40 seconds each. In the visual avoidance test, a black dot is directly controllable via a computer mouse. The operator moved the dot toward the tadpole. If the tadpole responded, the operator would proceed to initiate the next encounter. If the tadpole did not respond, the operator would initiate a new encounter after five seconds. In the edge preference test, the entire screen is white. The setup is placed inside a dark box to minimize light contamination. The interior of the box is coated black to minimize reflections from the screen. Tadpoles are recorded using a video camera pointed down on the tank through a hole in the top of the box. **b-i,** Time-lapse snapshots of a visual avoidance video showing the trajectory process of a behaving tadpole. The colored lines connect the position of the tadpole in adjacent frames to form a trajectory. Crosses are labeled in frames where the tadpole met the black dot. **j-l,** Representative traces of behavior test data (top) and corresponding analyzed data (bottom). **j,** (Top) representative trajectories of tadpole movement in a color preference test. The green dotted lines indicate the boundary between the black and white areas. The right and left areas are white and black from 0-40 seconds, and switch colors from 40-80 seconds. Dashed and solid lines represent the trajectories of tadpole movement from 0-40 seconds and 40-80 seconds, respectively. (Bottom) distance of the tadpoles to the black and white boundary. Dashed and solid lines represent the trajectories of tadpole movement from 0-40 seconds and 40-80 seconds, respectively. Gray color highlights the black side. **k,** (Top) representative trajectories of tadpole movement in a visual avoidance test. Green crosses indicate the locations where the tadpole encountered the black dots. (Bottom) distance between the tadpole and the black dot during the test. **l,** (Top) representative trajectories of tadpoles in an edge preference test. The green ring indicates the outer quarter radius of the container defined as the edge in the experiment. (Bottom) distance between the tadpole and the container center during the test. The green color indicates the edge region. The statistic results of behavior tests, including examples in (**j-l**), were presented in Fig. 3o**-q**.

**Extended Data Fig. 7.**
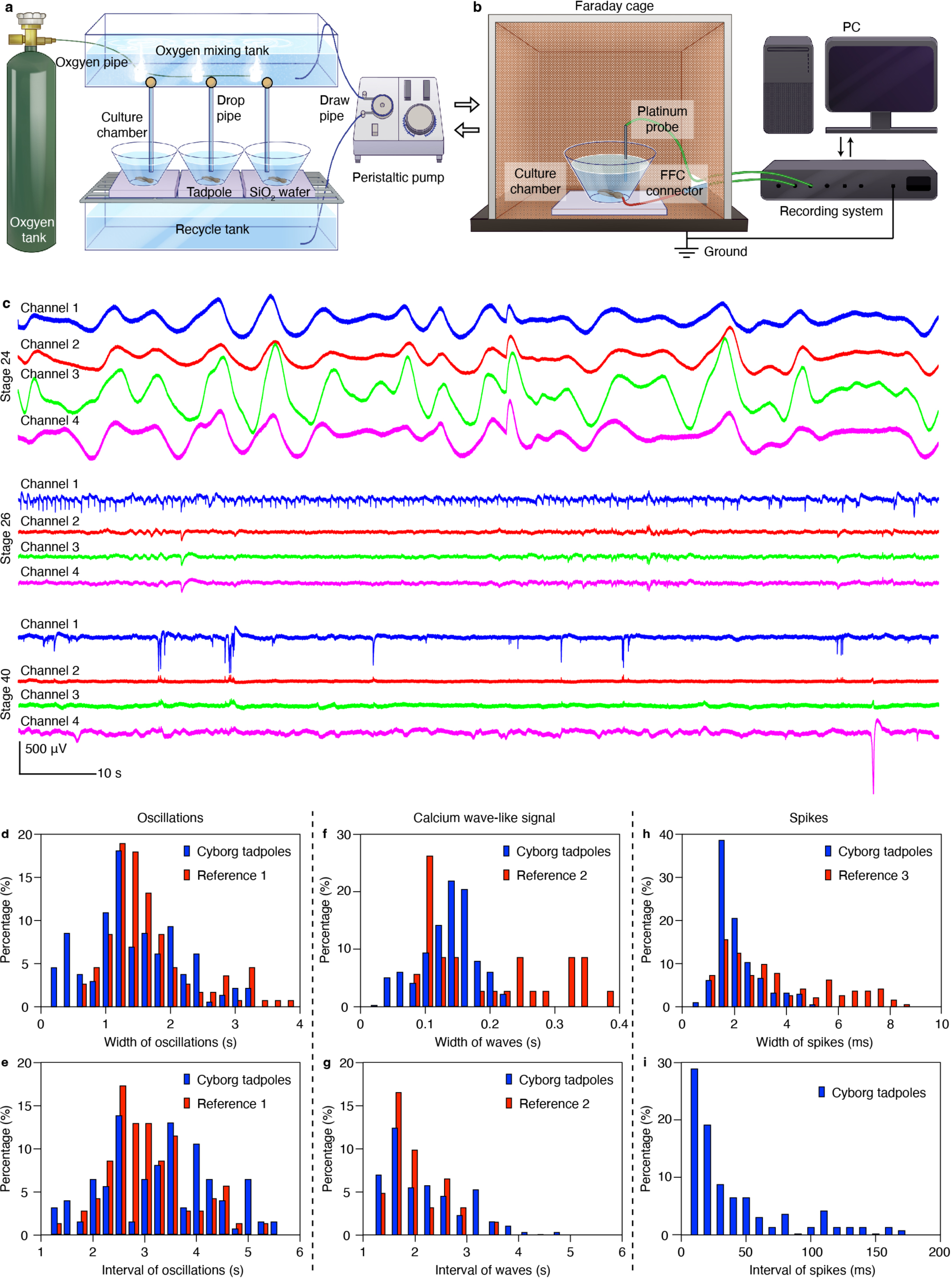
Experimental setup, raw data, and reference comparison of continuous electrophysiology in *Xenopus* embryonic brain development. **a,** Schematics showing the oxygen anesthetic system used to minimize tadpole movement during culture for recording. The system mixes the anesthetic media with fresh oxygen as previously reported^40^ to minimize the effects of anesthesia on tadpole development. **b,** Schematics showing the recording setup for electrophysiological experiments. During recording, the culture chamber is placed in a Faraday cage on a grounded optic table. The input/output of the implanted mesh electronics is connected to a recording system using a flexible flat cable (FFC) connector. A Pt probe is placed in the culture media as ground. **c,** Raw data of continuous recordings shown in (Fig. 4). **d-i,** Reference comparison of continuous electrophysiology. Distribution plots showing comparisons of oscillation signal width (**d**) and interval (**e**); calcium-wave like signal width (**f**) and interval (**g**); spike width (**h**) and interval (**i**). Reference data is as follows: reference 1^59^, reference 2^41^, and reference 3^42^. Reference 3 did not include the corresponding dataset for spike intervals, so it is not included in (**i**). The results in (**d-i**) are determined from signals collected from three cyborg tadpoles.

**Extended Data Fig. 8.**
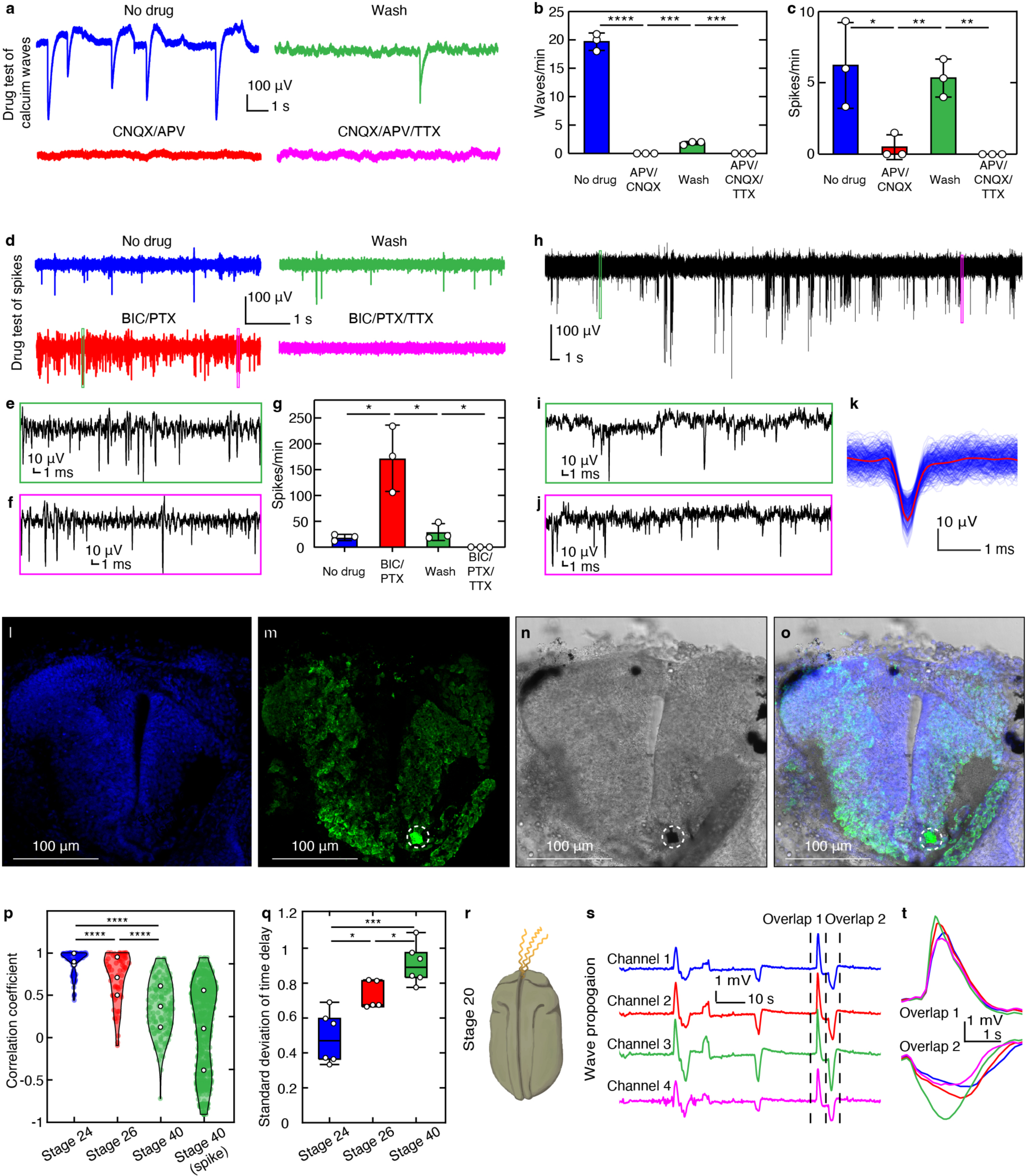
Analysis of continuous electrophysiology in *Xenopus* embryonic brain development. **a-g,** Pharmacological tests. **a,** Dynamics of calcium wave-like signals during the pharmacological test. Representative voltage traces from the cyborg tadpole under serial pharmacological test conditions of no drug, cyanquixaline (CNQX)/[2R]-amino-5-phosphonopentanoate (APV), wash of CNQX/APV, and CNQX/APV/ tetrodotoxin (TTX). **b,** The change of wave number per minute during the pharmacological test. Bar plots indicate mean ± s.d., each dot represents a recording trial. Two-tailed unpaired t-test, ***, *p* < 0.001, ****, *p* < 0.0001. **c-g,** Dynamics of spike-like signals during the pharmacological test. **c,** Statistical summary of the firing rate changes under different conditions. The tadpole was treated with APV/CNQX, washed, then CNQX/APV/ TTX in series. Bar plots indicate mean ± s.d., each dot represents a recording trial. Two-tailed unpaired t-test, *, *p* < 0.05, **, *p* < 0.01. **d,** Representative voltage traces from the cyborg tadpole under serial pharmacological test conditions of no drug, bicuculline (BIC)/picrotoxin (PTX), washed, followed by BIC/PTX/TTX. **e, f,** Zoomed-in views of the signal highlighted by the green (**e**) and magenta (**f**) boxes in (**d**). **g,** Statistical summary of the firing rate changes under different conditions. The tadpole was treated with BIC/PTX, washed, then BIC/PTX/TTX in series. Bar plots indicate mean ± s.d., each dot represents a recording trial. Two-tailed unpaired t-test, *, *p* < 0.05. **h-o,** Correlation of single-unit action potential signals with the corresponded electrode position. **h,** Representative voltage traces from a cyborg tadpole showing single-unit action potentials. **i, j,** Zoomed-in views of the signal highlighted by green (**i**) and magenta (**j**) boxes in (**h**). **k,** Mean spike superimposed on all spikes from the same unit, sorted by spike sorting of data in (**h**). **l-o,** Confocal fluorescence images of the cyborg tadpole brain slice showing DAPI (**l**), HuC/D (**m**), BF (**n**) and overlaid (**o**) channels. The white dashed circles highlight the position of the electrode that recorded the voltage trace in (**h**). **p,** Correlation coefficient between channels of stages 24, 26, stage 40 local field potential and stage 40 spike signals. Positive correlation corresponds to a coefficient of 1, negative to -1, and no correlation to 0. White dots represent the lower quartile, median, upper quartile from bottom to top. Each translucent dot represents a sample. Two-tailed unpaired t-test, ****, *p* < 0.0001. **q,** Standard deviation of time delay between channels of stages 24, 26, and 40 local field potential signals. Lower standard deviations indicate greater synchronization between channels. Box plots indicate minimum, lower quartile, median, upper quartile, and maximum. Each white dot represents a sample. Two-tailed unpaired t-test, *, *p* < 0.05, ***, *p* < 0.001. **r-t,** Propagating wave signals in stage 20 embryonic brain. **r,** Schematic of the cyborg tadpole at developmental stage 20. **s,** Representative voltage traces from four channels in the cyborg tadpole at stage 20. **t,** Zoomed-in views of the signals highlighted by dashed lines in (**s**).

**Extended Data Fig. 9.**
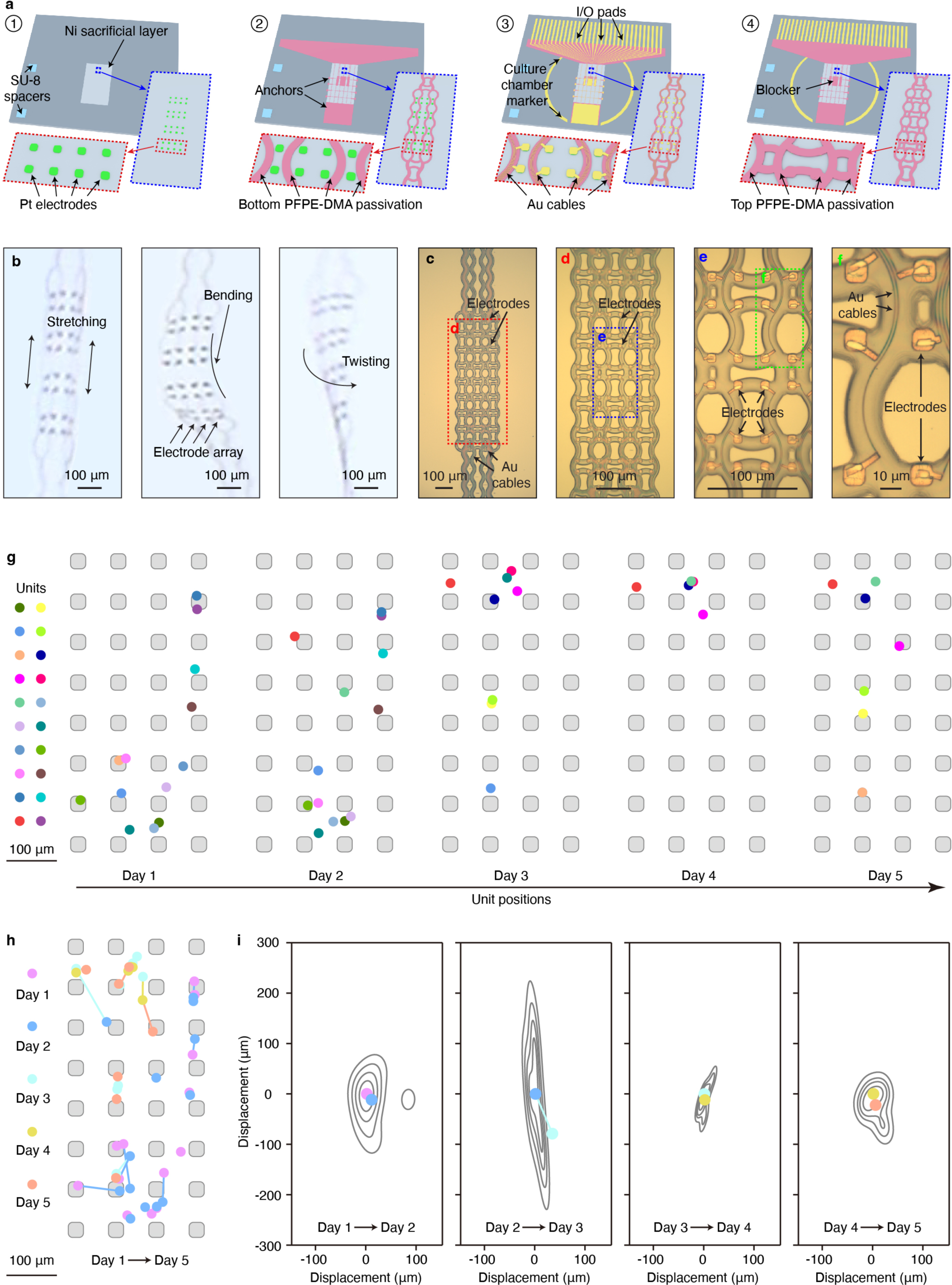
Soft and stretchable high-density mesh electrode array for tracking neural activities. **a,** Schematics showing the electron-beam (e-beam) lithographic fabrication of PFPE-DMA-encapsulated stretchable mesh electronics with a 32-channel high-density electrode array. First, a Ni layer is deposited on a blank silicon oxide wafer as a sacrificial layer. A SU-8 layer is patterned as a spacer, Pt electrodes are photolithographically patterned (step 1). Then, the bottom PFPE-DMA (step 2), Au interconnects (step 3), and top PFPE-DMA layer (step 4) are lithographically patterned. Au layer is patterned by e-beam lithography. PFPE-DMA layers are patterned by photolithography. Zoomed-in images show the details of the electrode arrays (highlighted in blue dashed boxes) and individual electrodes (highlighted in red dashed boxes). **b,** Photographic images showing the free-floating 32-channel high-density electrode array during stretching, bending, and twisting. **c,** BF microscopic image showing a representative 128-channel electrode array. **d,** Zoomed-in view of the red dashed box-highlighted region in (**c**) showing the high-density electrodes and interconnects. **e,** Zoomed-in view of the blue dashed box-highlighted region in (**d**) showing the stretchable design. **f,** Zoomed-in view of the green dashed box-highlighted region in (**e**) showing the individual electrodes and interconnects. **g,** Single-unit waveform centroids (*n* = 20 neurons) from a continuous 5-day recording in the cyborg axolotl tadpole (centroid computed using spatial average across electrode positions weighted by the square of the mean waveform amplitude at each electrode). Grey patterns indicate the positions and sizes of the mesh electrodes. **h,** Single-neuron waveform centroid displacement throughout the 5-day recording in the cyborg axolotl tadpole. Centroids from the same day are labeled with the same color. Centroids for the same neurons across different days are connected by lines. Grey patterns indicate the positions and sizes of the mesh electrodes. **i,** Average displacement of single-neuron centroids across different days. Grey contours indicate quintile boundaries of the distribution of centroid position displacement.

**Extended Data Fig. 10.**
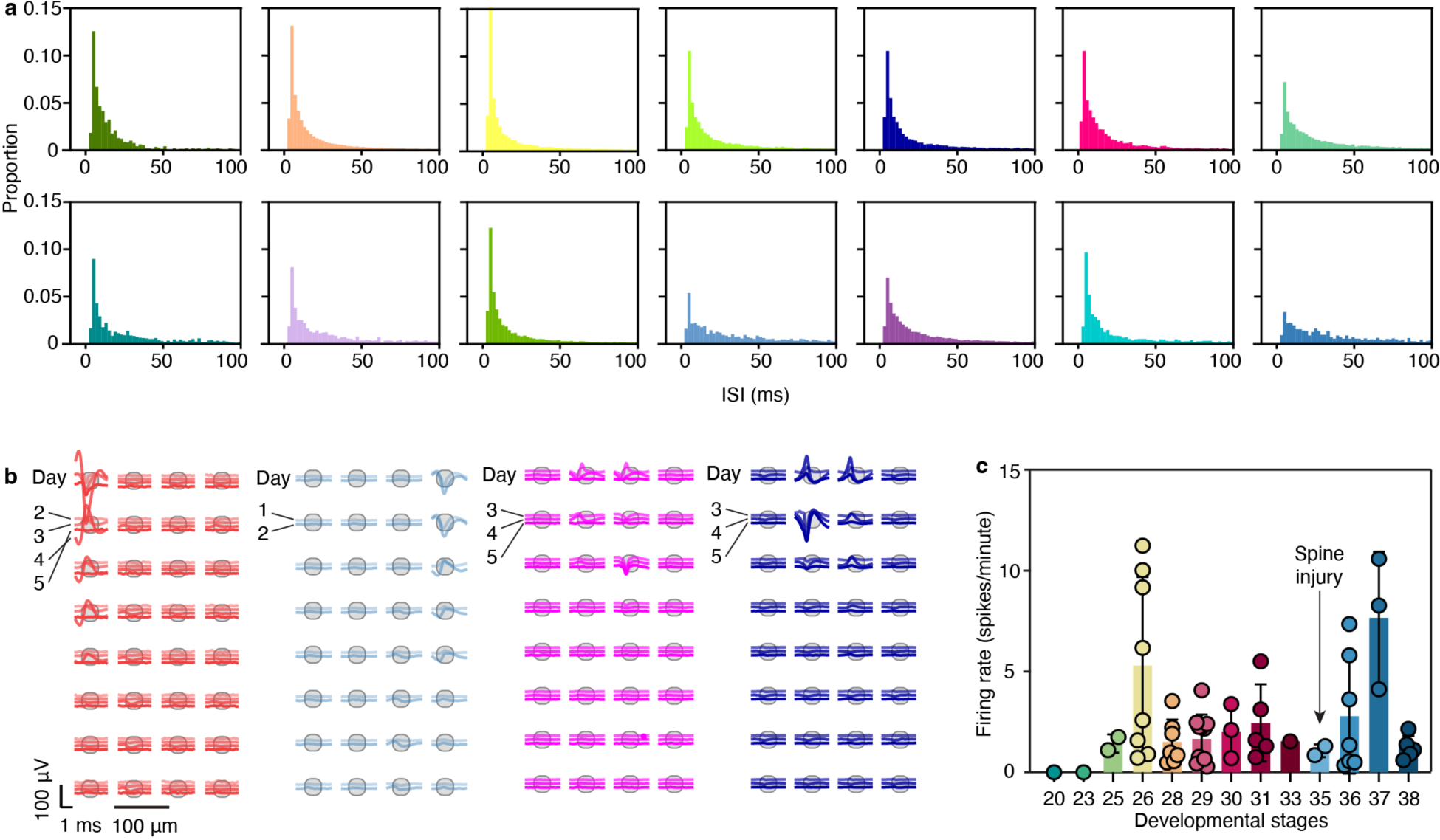
Analysis of continuous electrophysiology in axolotl embryonic brain development. **a,** Representative inter-spike interval (ISI) of spikes sorted from axolotl recording. Each color corresponds to an identified unit. **b,** Representative average single-unit waveforms at each of the recording electrodes over the 5-day of recording in a cyborg axolotl tadpole. **c,** Time evolution of the spike firing rate in cyborg axolotl embryo development and spinal cord injury regeneration. Bar plots indicate mean ± s.d., each dot represents the firing rate of one unit.

## Supplementary information

The supplementary information document contains Supplementary discussion, Supplementary Figures 1-8 and Supplementary Tables 1-3.

## Supplementary discussion

### 1. Face-down electrodes

During implantation, the mesh needs to be attached to an embryo, with its bottom side in contact with the neural plate (**Supplementary Fig. 1a**). Therefore, to enable signal recording, the electrodes must be exposed on the bottom side of the electronics. We developed a fabrication recipe to expose the electrodes on the bottom surface of the mesh, ensuring direct contact with the neural plate.

### 2. I/O pad design

Conventional SU-8 electronics utilized SU-8 as a back layer for the I/O pad. However, PFPE-DMA, being too soft, would beak during flip-chip bonding if still used as the back layer. Therefore, we designed the I/O pads without the bottom passivation (**Supplementary Fig. 1b**).

### 3. Choosing PFPE-DMA with a molecular weight of 8 kDa

We chose the 8 kDa PFPE-DMA based on the fabrication yield. To increase the adhesion between the Au layer and the PFPE-DMA layer during Au deposition. We developed a new protocol that treats the PFPE-DMA surface with inert gas plasma, and then uses sputtering to deposit the Au layer. However, the sputtered Au layer was difficult to lift off. Our findings showed that a device made of softer PFPE-DMA had a lower overall lift-off yield, with yields of PFPE-DMA with molecular weights of 4-, 8-, 10-, and 12-kDa being around 95%, 90%, 70%, and 30%, respectively. As the result, we decided to choose 8 kDa PFPE-DMA, which has both decent softness and fabrication yield.

**Supplementary Fig. 1.**
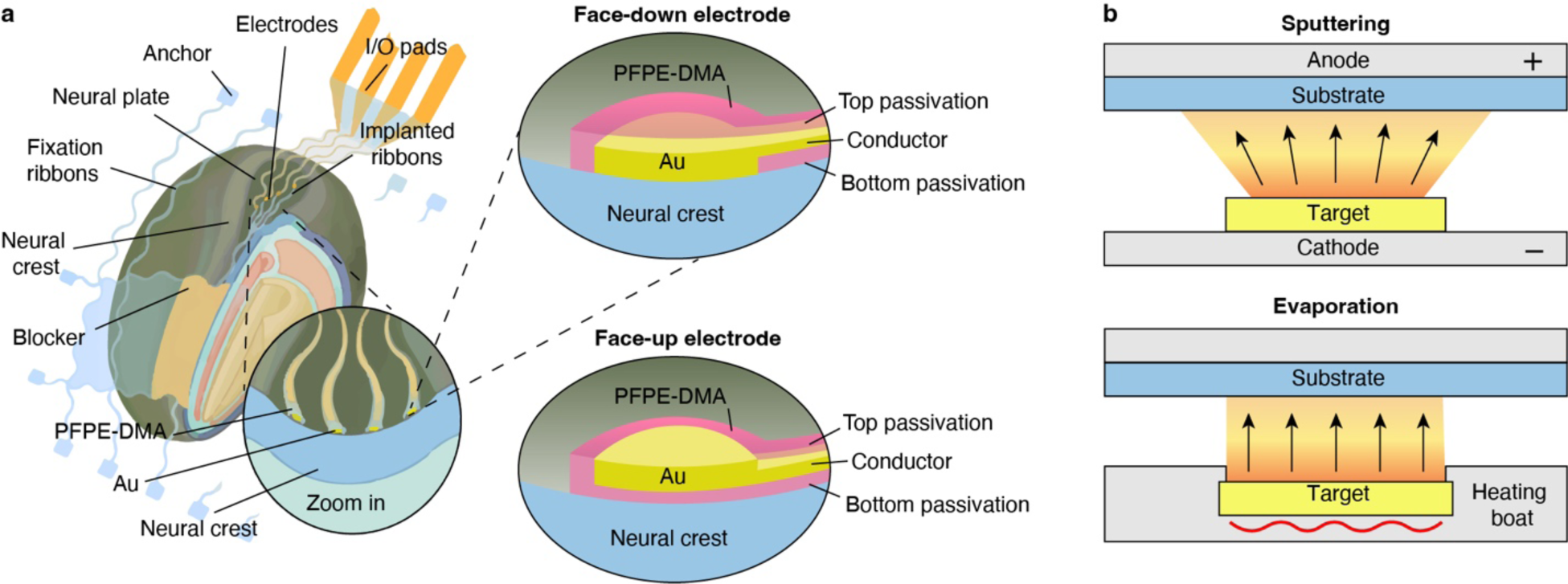
Technical innovations critical to soft and stretchable PFPE-DMA mesh electronics. **a,** Schematics comparing face-up and face-down electrodes in soft and stretchable mesh electronics for embryo implantation and neural interfaces. **b,** Schematics showing sputtering (left) and evaporation (right) for metal deposition on PFPE-DMA layers.

**Supplementary Fig. 2.**
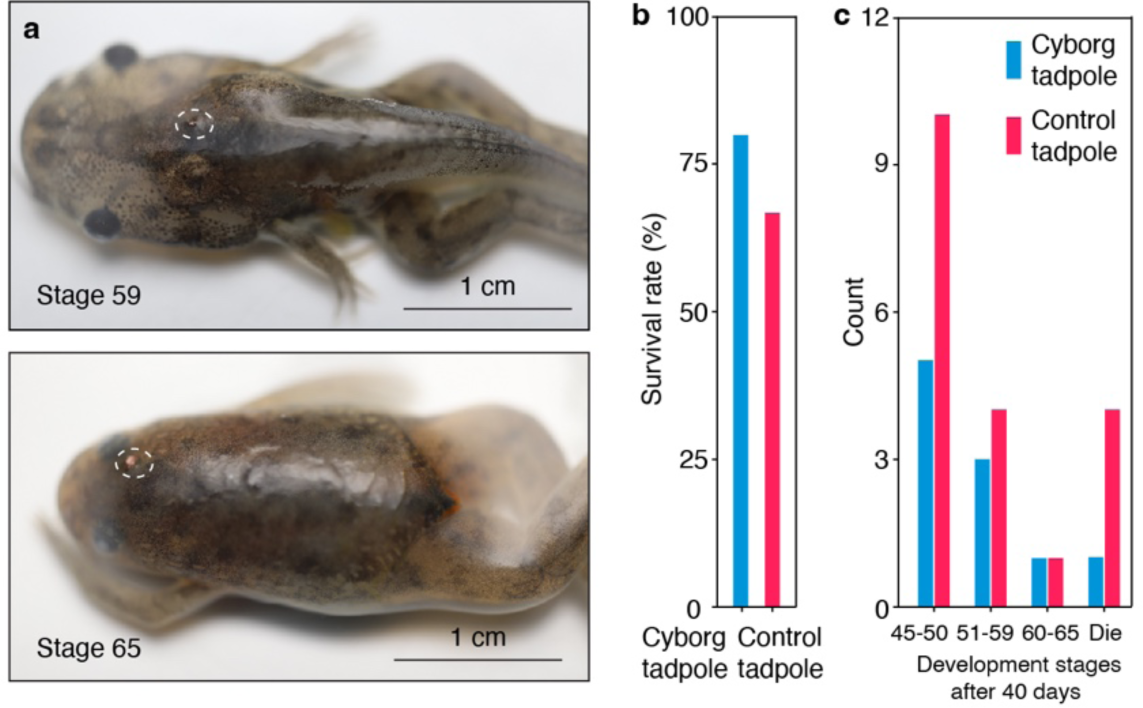
Long-term rearing of cyborg tadpoles to cyborg frogs. **a,** Photos showing cyborg frogs at stage 59 (top) and stage 65 (bottom). Dashed circles highlight the interconnects of mesh electronics outside the brain. **b,** Survival rate of cyborg and control tadpoles after 40 days of rearing. **c,** Development stages of cyborg and control tadpoles after 40 days of rearing.

**Supplementary Fig. 3.**
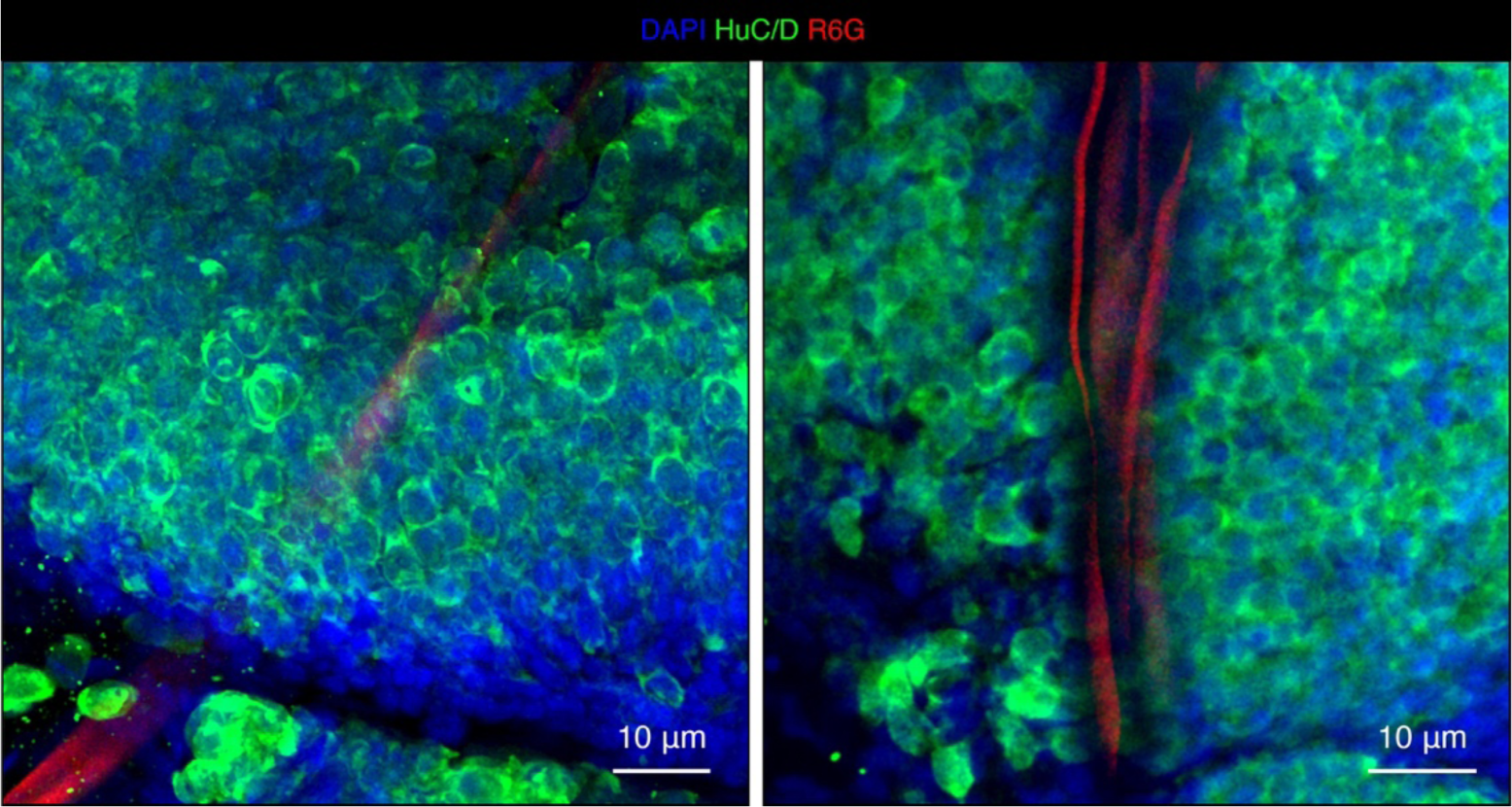
Immunostaining images depicting the contact between *Xenopus* brain tissues and mesh electronics. Whole-mount-stained 3D reconstructed confocal fluorescence images of implanted mesh microelectronics showing the mesh embedded in the neural tissue. 4′,6-diamidino-2-phenylindole (DAPI) labels cell nuclei, HuC/D labels neurons, and Rhodamine 6G (R6G) labels the device.

**Supplementary Table 1.**
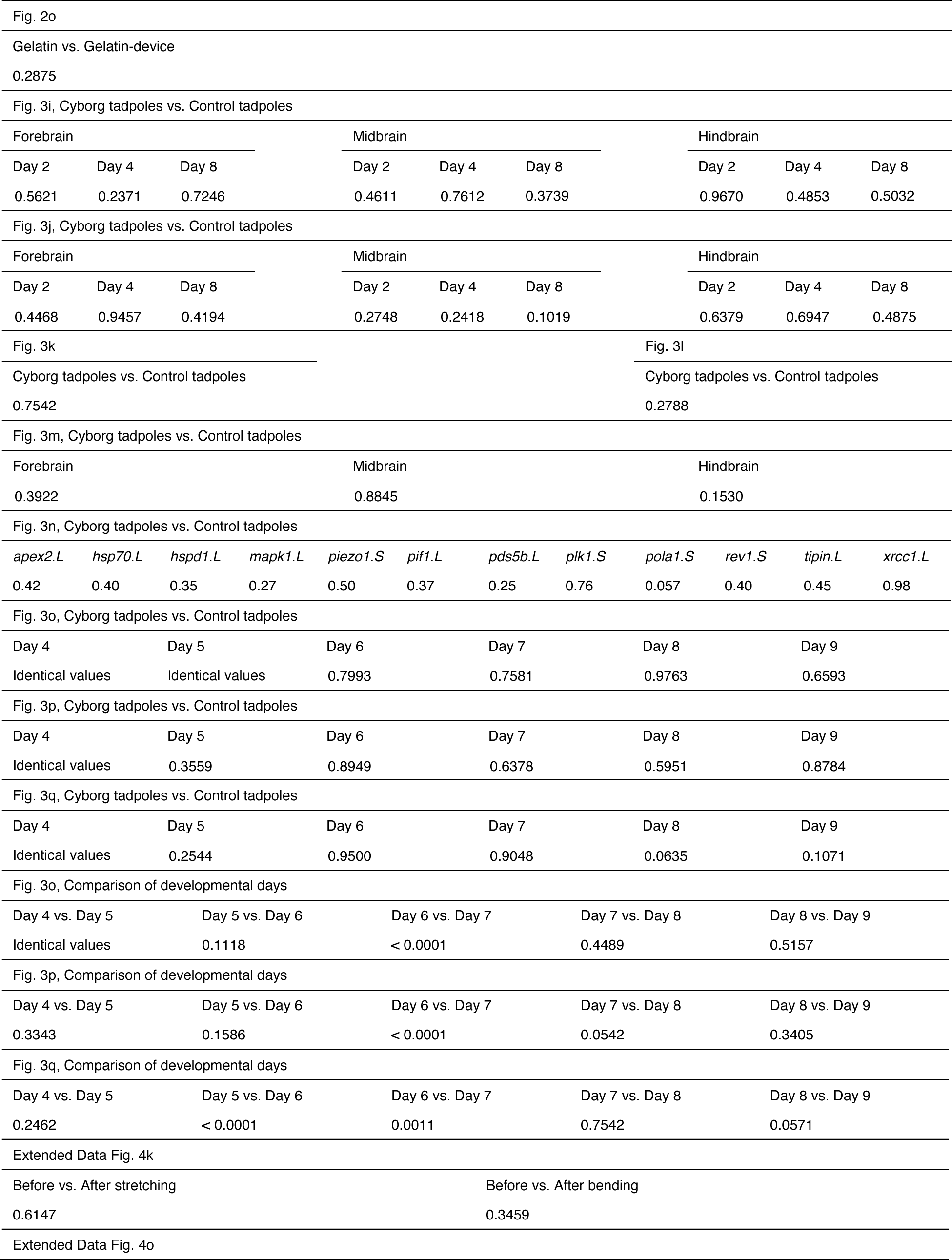
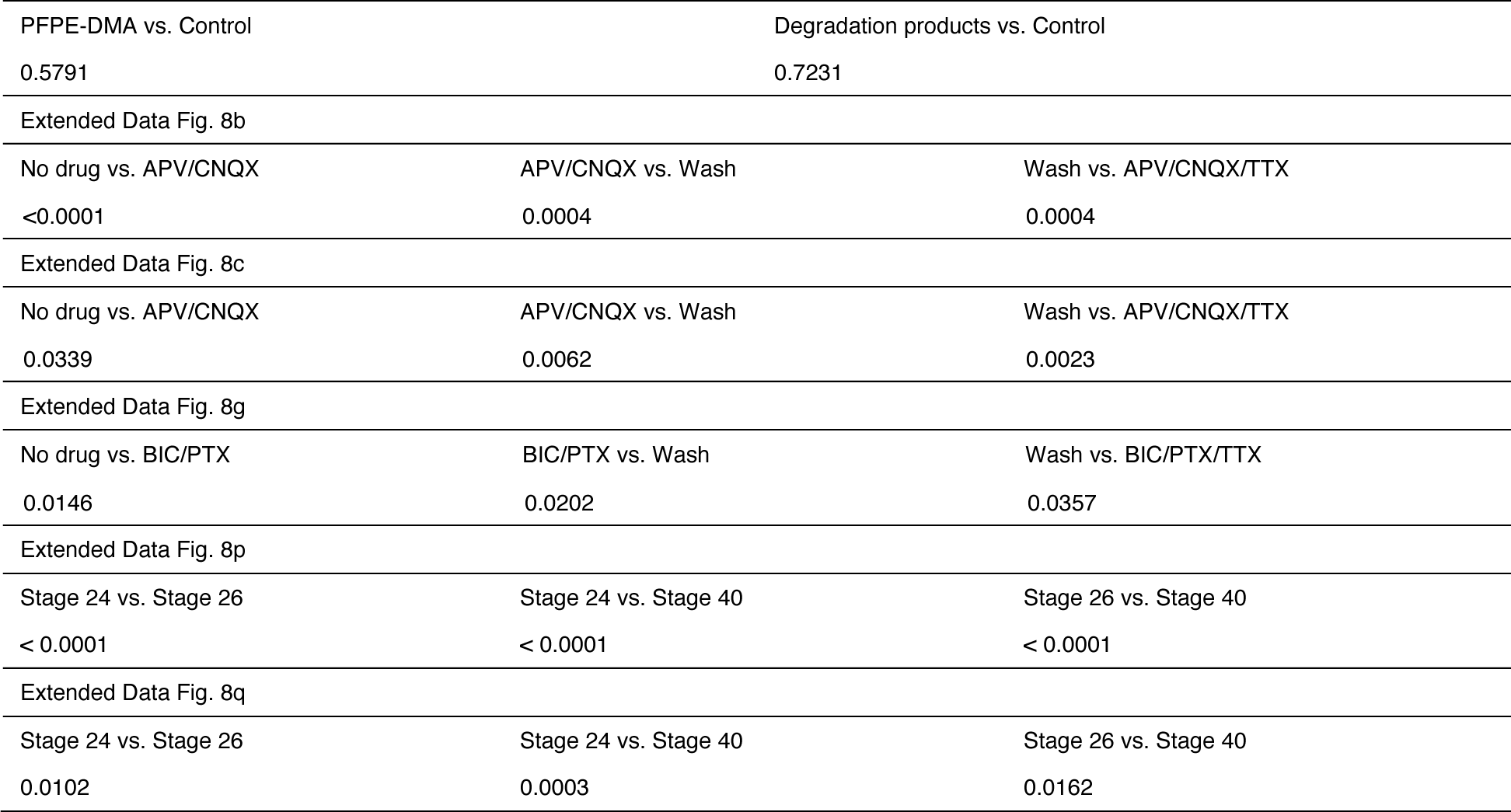
Actual p values in two-tailed unpaired t-tests.

|  |  |  |  |  |  |  |  |  |  |  |  |
| --- | --- | --- | --- | --- | --- | --- | --- | --- | --- | --- | --- |
| Fig. 2o |  |  |  |  |  |  |  |  |  |  |  |
| Gelatin vs. Gelatin-device |  |  |  |  |  |  |  |  |  |  |  |
| 0.2875 |  |  |  |  |  |  |  |  |  |  |  |
| Fig. 3i, Cyborg tadpoles vs. Control tadpoles |  |  |  |  |  |  |  |  |  |  |  |
| Forebrain |  |  | Midbrain |  |  | Hindbrain |  |  |  |  |  |
| Day 2 | Day 4 | Day 8 | Day 2 | Day 4 | Day 8 | Day 2 | Day 4 | Day 8 |  |  |  |
| 0.5621 | 0.2371 | 0.7246 | 0.4611 | 0.7612 | 0.3739 | 0.9670 | 0.4853 | 0.5032 |  |  |  |
| Fig. 3j, Cyborg tadpoles vs. Control tadpoles |  |  |  |  |  |  |  |  |  |  |  |
| Forebrain |  |  | Midbrain |  |  | Hindbrain |  |  |  |  |  |
| Day 2 | Day 4 | Day 8 | Day 2 | Day 4 | Day 8 | Day 2 | Day 4 | Day 8 |  |  |  |
| 0.4468 | 0.9457 | 0.4194 | 0.2748 | 0.2418 | 0.1019 | 0.6379 | 0.6947 | 0.4875 |  |  |  |
| Fig. 3k |  |  |  |  |  | Fig. 3l |  |  |  |  |  |
| Cyborg tadpoles vs. Control tadpoles |  |  |  |  |  | Cyborg tadpoles vs. Control tadpoles |  |  |  |  |  |
| 0.7542 |  |  |  |  |  | 0.2788 |  |  |  |  |  |
| Fig. 3m, Cyborg tadpoles vs. Control tadpoles |  |  |  |  |  |  |  |  |  |  |  |
| Forebrain |  |  | Midbrain |  |  | Hindbrain |  |  |  |  |  |
| 0.3922 |  |  | 0.8845 |  |  | 0.1530 |  |  |  |  |  |
| Fig. 3n, Cyborg tadpoles vs. Control tadpoles |  |  |  |  |  |  |  |  |  |  |  |
| *apex2.L* | *hsp70.L* | *hspd1.L* | *mapk1.L* | *piezo1.S* | *pif1.L* | *pds5b.L* | *plk1.S* | *pola1.S* | *rev1.S* | *tipin.L* | *xrcc1.L* |
| 0.42 | 0.40 | 0.35 | 0.27 | 0.50 | 0.37 | 0.25 | 0.76 | 0.057 | 0.40 | 0.45 | 0.98 |
| Fig. 3o, Cyborg tadpoles vs. Control tadpoles |  |  |  |  |  |  |  |  |  |  |  |
| Day 4 |  | Day 5 |  | Day 6 |  | Day 7 |  | Day 8 |  | Day 9 |  |
| Identical values |  | Identical values |  | 0.7993 |  | 0.7581 |  | 0.9763 |  | 0.6593 |  |
| Fig. 3p, Cyborg tadpoles vs. Control tadpoles |  |  |  |  |  |  |  |  |  |  |  |
| Day 4 |  | Day 5 |  | Day 6 |  | Day 7 |  | Day 8 |  | Day 9 |  |
| Identical values |  | 0.3559 |  | 0.8949 |  | 0.6378 |  | 0.5951 |  | 0.8784 |  |
| Fig. 3q, Cyborg tadpoles vs. Control tadpoles |  |  |  |  |  |  |  |  |  |  |  |
| Day 4 |  | Day 5 |  | Day 6 |  | Day 7 |  | Day 8 |  | Day 9 |  |
| Identical values |  | 0.2544 |  | 0.9500 |  | 0.9048 |  | 0.0635 |  | 0.1071 |  |
| Fig. 3o, Comparison of developmental days |  |  |  |  |  |  |  |  |  |  |  |
| Day 4 vs. Day 5 |  | Day 5 vs. Day 6 |  | Day 6 vs. Day 7 |  | Day 7 vs. Day 8 |  | Day 8 vs. Day 9 |  |  |  |
| Identical values |  | 0.1118 |  | < 0.0001 |  | 0.4489 |  | 0.5157 |  |  |  |
| Fig. 3p, Comparison of developmental days |  |  |  |  |  |  |  |  |  |  |  |
| Day 4 vs. Day 5 |  | Day 5 vs. Day 6 |  | Day 6 vs. Day 7 |  | Day 7 vs. Day 8 |  | Day 8 vs. Day 9 |  |  |  |
| 0.3343 |  | 0.1586 |  | < 0.0001 |  | 0.0542 |  | 0.3405 |  |  |  |
| Fig. 3q, Comparison of developmental days |  |  |  |  |  |  |  |  |  |  |  |
| Day 4 vs. Day 5 |  | Day 5 vs. Day 6 |  | Day 6 vs. Day 7 |  | Day 7 vs. Day 8 |  | Day 8 vs. Day 9 |  |  |  |
| 0.2462 |  | < 0.0001 |  | 0.0011 |  | 0.7542 |  | 0.0571 |  |  |  |
| Extended Data Fig. 4k |  |  |  |  |  |  |  |  |  |  |  |
| Before vs. After stretching |  |  |  |  |  | Before vs. After bending |  |  |  |  |  |
| 0.6147 |  |  |  |  |  | 0.3459 |  |  |  |  |  |
| Extended Data Fig. 4o |  |  |  |  |  |  |  |  |  |  |  |

|  |  |  |
| --- | --- | --- |
| PFPE-DMA vs. Control |  | Degradation products vs. Control |
| 0.5791 |  | 0.7231 |
| Extended Data Fig. 8b |  |  |
| No drug vs. APV/CNQX | APV/CNQX vs. Wash | Wash vs. APV/CNQX/TTX |
| <0.0001 | 0.0004 | 0.0004 |
| Extended Data Fig. 8c |  |  |
| No drug vs. APV/CNQX | APV/CNQX vs. Wash | Wash vs. APV/CNQX/TTX |
| 0.0339 | 0.0062 | 0.0023 |
| Extended Data Fig. 8g |  |  |
| No drug vs. BIC/PTX | BIC/PTX vs. Wash | Wash vs. BIC/PTX/TTX |
| 0.0146 | 0.0202 | 0.0357 |
| Extended Data Fig. 8p |  |  |
| Stage 24 vs. Stage 26 | Stage 24 vs. Stage 40 | Stage 26 vs. Stage 40 |
| < 0.0001 | < 0.0001 | < 0.0001 |
| Extended Data Fig. 8q |  |  |
| Stage 24 vs. Stage 26 | Stage 24 vs. Stage 40 | Stage 26 vs. Stage 40 |
| 0.0102 | 0.0003 | 0.0162 |

**Supplementary Table 2.** Primary and secondary antibodies.

| Name | Vendor | Catalog Number | Concentration |
| --- | --- | --- | --- |
| Recombinant Anti-HuD + HuC antibody | Abcam | AB184267 | 1:100 |
| Anti-Acetylated Tubulin antibody, Mouse monoclonal | Sigma-Aldrich | T7451 | 1:100 |
| BrdU Monoclonal Antibody (MoBU-1) | Invitrogen | B35128 | 1:100 |
| Anti-SOX2 Monoclonal (Btjce) | Invitrogen | 14-9811-82 | 1:100 |
| Anti-Myt1/MTF1 antibody | Abcam | AB251682 | 1:100 |
| Anti-vimentin antibody (SP20) | Abcam | AB16700 | 1:100 |
| Ms mAb to ALDH1L1 (3E9) | Abcam | AB56777 | 1:100 |
| Goat anti-Rat IgG (H+L) Cross-Adsorbed Secondary Antibody, Alexa Fluor™ 488 | Invitrogen | A-11006 | 1:500 |
| Goat anti-Rabbit IgG (H+L) Cross-Adsorbed Secondary Antibody, Alexa Fluor™ 594 | Invitrogen | A-11012 | 1:500 |
| IgG (H+L) Highly Cross-Adsorbed Donkey anti-Mouse, Alexa Fluor™ Plus 647 | Invitrogen | A-32787 | 1:500 |
| Goat anti-Rabbit IgG (H+L) Highly Cross-Adsorbed Secondary Antibody, Alexa Fluor™ Plus 488 | Invitrogen | A-32731 | 1:500 |
| Alexa Fluor™ 647 Phalloidin | Thermo Fisher<br>SCIENTIFIC | A22287 | 1:50 |

**Supplementary Fig. 4.**
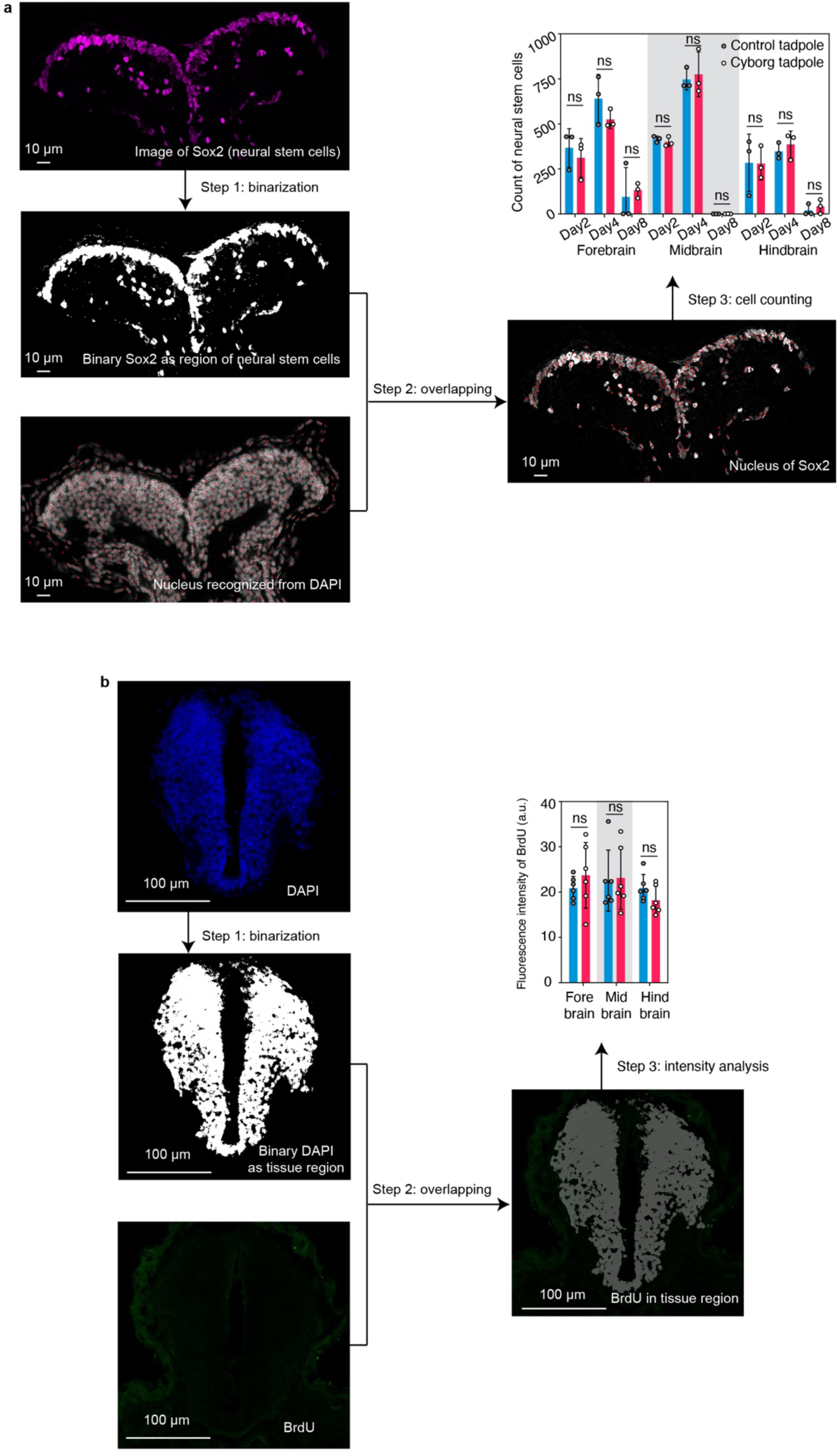
Procedures for quantitative analysis of fluorescence images. **a,** Cell counting in SRY-box transcription factor 2 (Sox2, neuron stem cells) fluorescence images. First, the Sox2 image is binarized to identify the region of neuron stem cells (step 1). Then, the binary Sox2 image is overlaid with the DAPI-labeled cell nuclei to indicate the nucleus of neuron stem cells (step 2). Finally, the number of neuron stem cell nuclei is counted and reported (step 3). **b,** Fluorescent intensity quantifying of bromodeoxyuridine (BrdU) images. First, the DAPI image is binarized to identify the tissue region (step 1). Then, the binary DAPI image is overlaid with the BrdU image (step 2). The fluorescent intensity of BrdU in the DAPI tissue region is calculated and reported (step 3).

**Supplementary Fig. 5.**
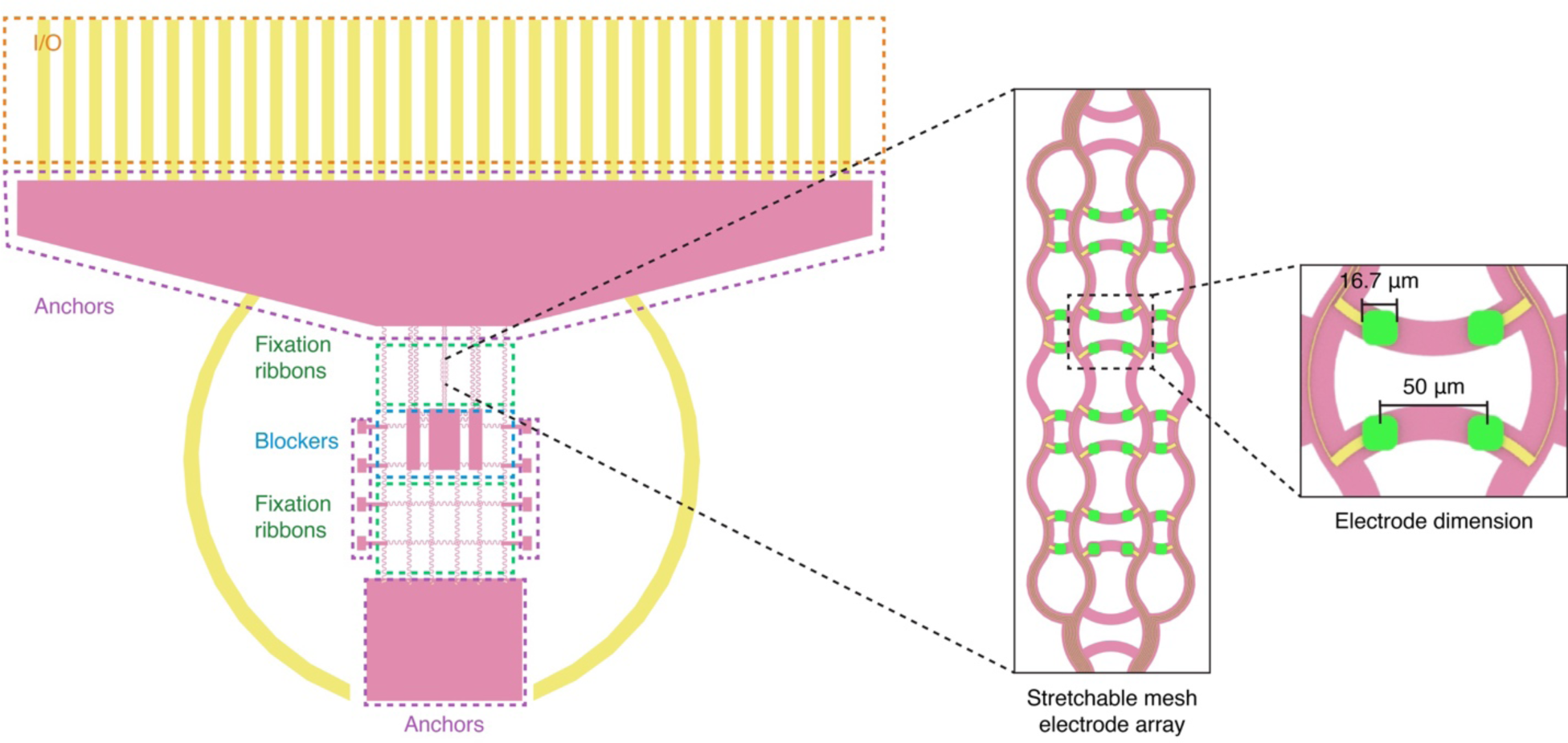
Design of the stretchable mesh electronics with a 32-channel mesh electrode array. The design contains a high-density mesh electrode array, stretchable anchors and ribbons, and blockers for embryo integration.

**Supplementary Table 3.**
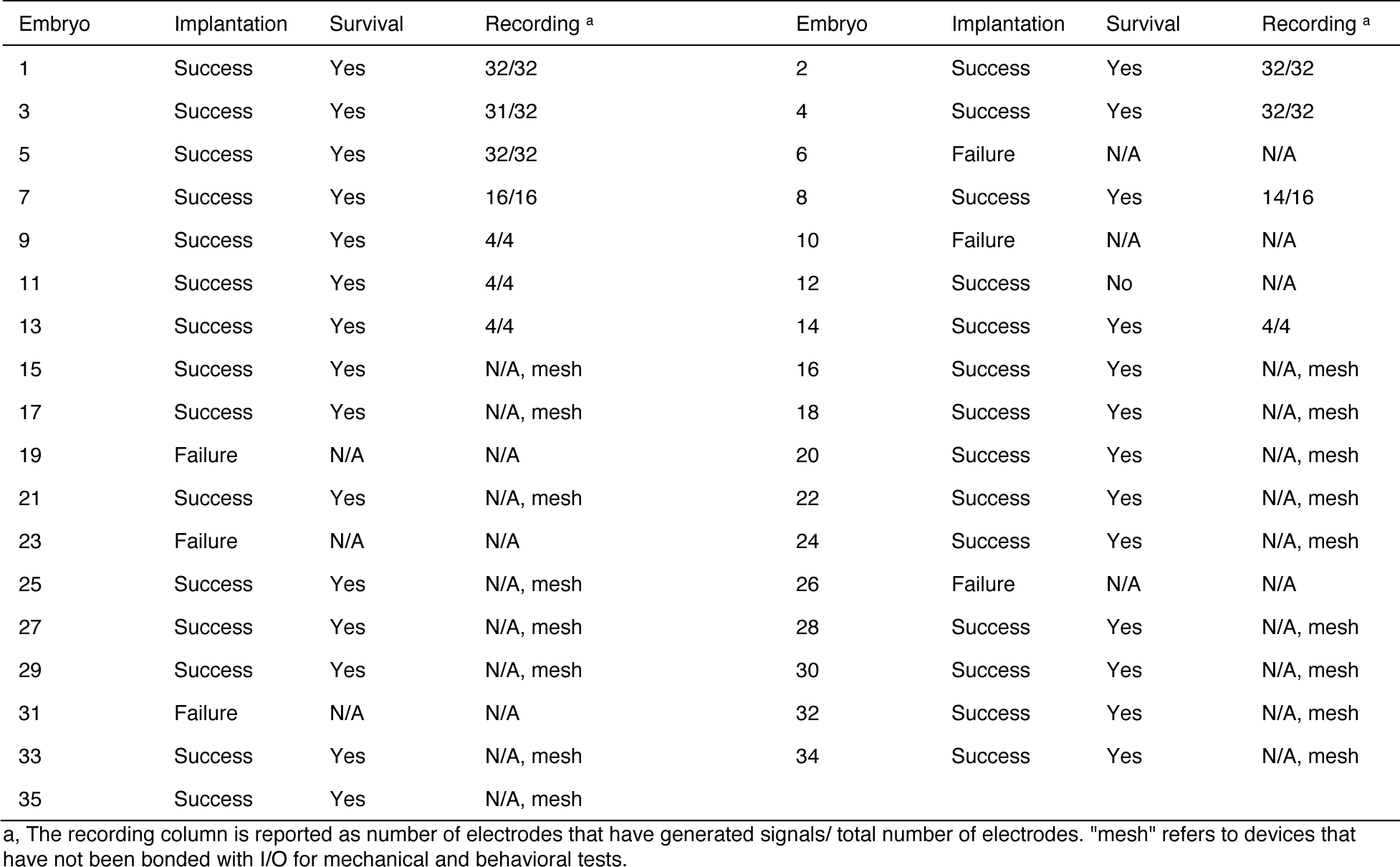
Yield of neurulation implantation.

**Supplementary Fig. 6.**
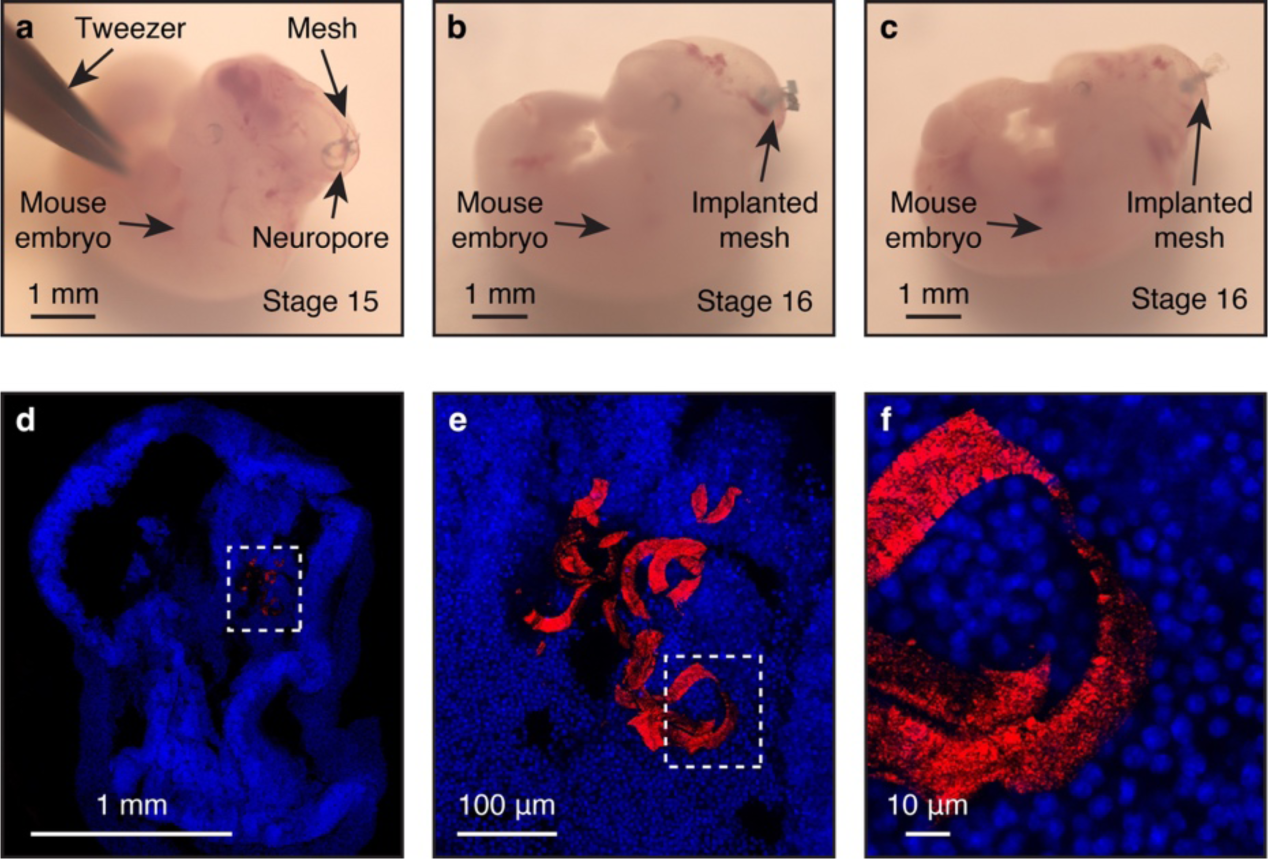
Implantation of stretchable mesh electronics in mouse embryos. **a-c,** BF microscopic images showing three representative mouse embryos implanted with stretchable mesh electronics. **d,** Confocal fluorescence image showing coronal sections of a cyborg mouse embryo fixed at embryonic stage 16. Cell nuclei, blue, and mesh electronics, red (reflective mode). **e,** Zoomed-in view of the white dashed box-highlighted region in (**d**) showing the 3D integration of mesh electronics with the brain. **f,** Zoomed-in view of the white dashed box-highlighted region in (**e**) showing one stretchable ribbon.

**Supplementary Fig. 7.**
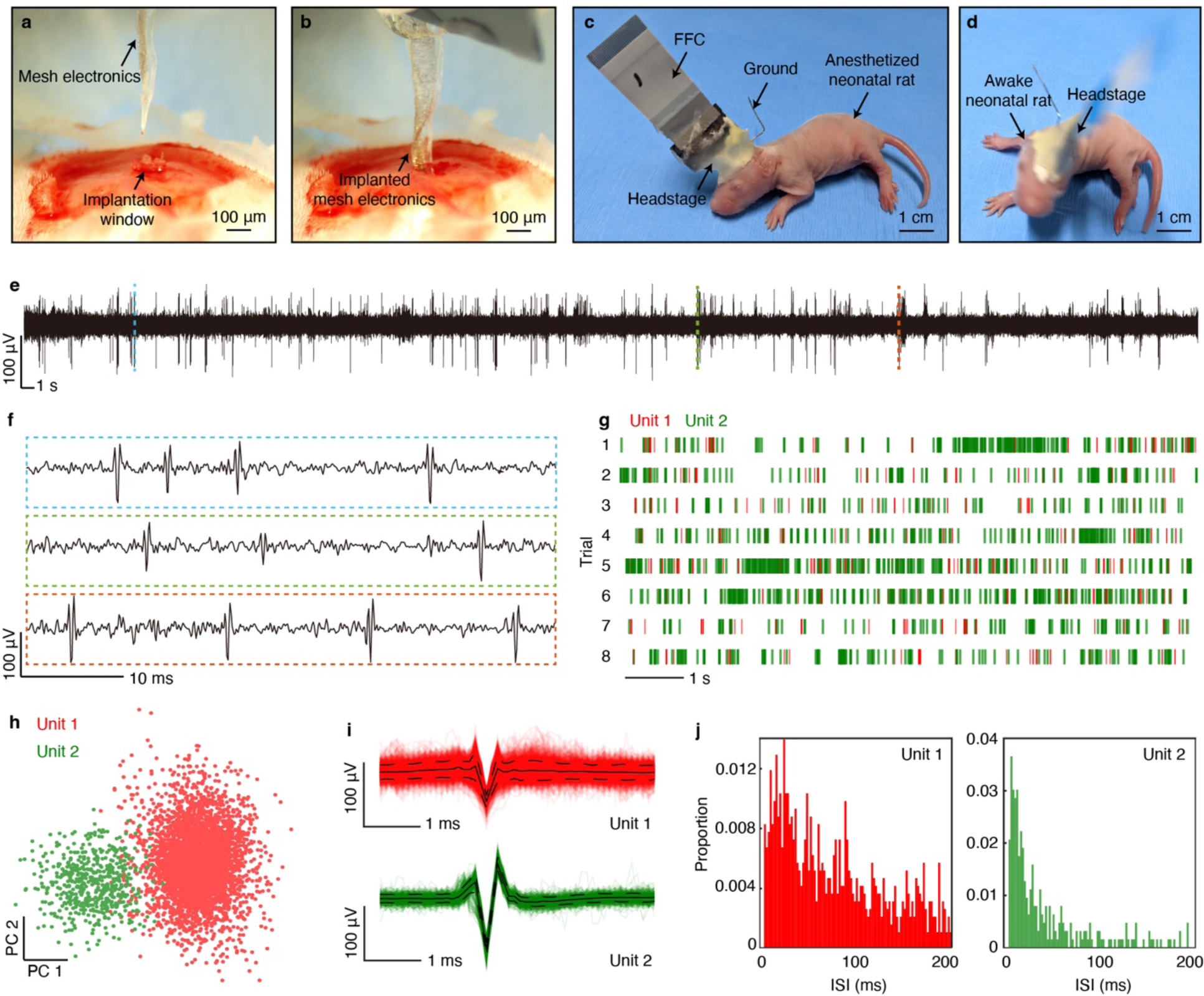
Implantation of stretchable mesh electronics in neonatal rat brain. **a, b,** Photographic images showing mesh electronics before (**a**) and after (**b**) implantation into a neonatal rat brain. **c, d,** Photographic images showing the neonatal rat after stereotactic surgery (**c**) and after recovering from anesthetics (**d**). **e,** Representative filtered voltage traces (300-3,000 Hz bandpass filter). **f,** Zoomed-in views of the signals highlighted by blue-, green- and red-dashed box-highlighted regions in (**e**). **g,** Raster plot of single-unit action potentials sorted from the recording. **h-j,** Principal component analysis (PCA) (**h**), average waveforms (mean ± s.d.) (**i**) and ISI (**j**) of two representative units.

**Supplementary Fig. 8.**
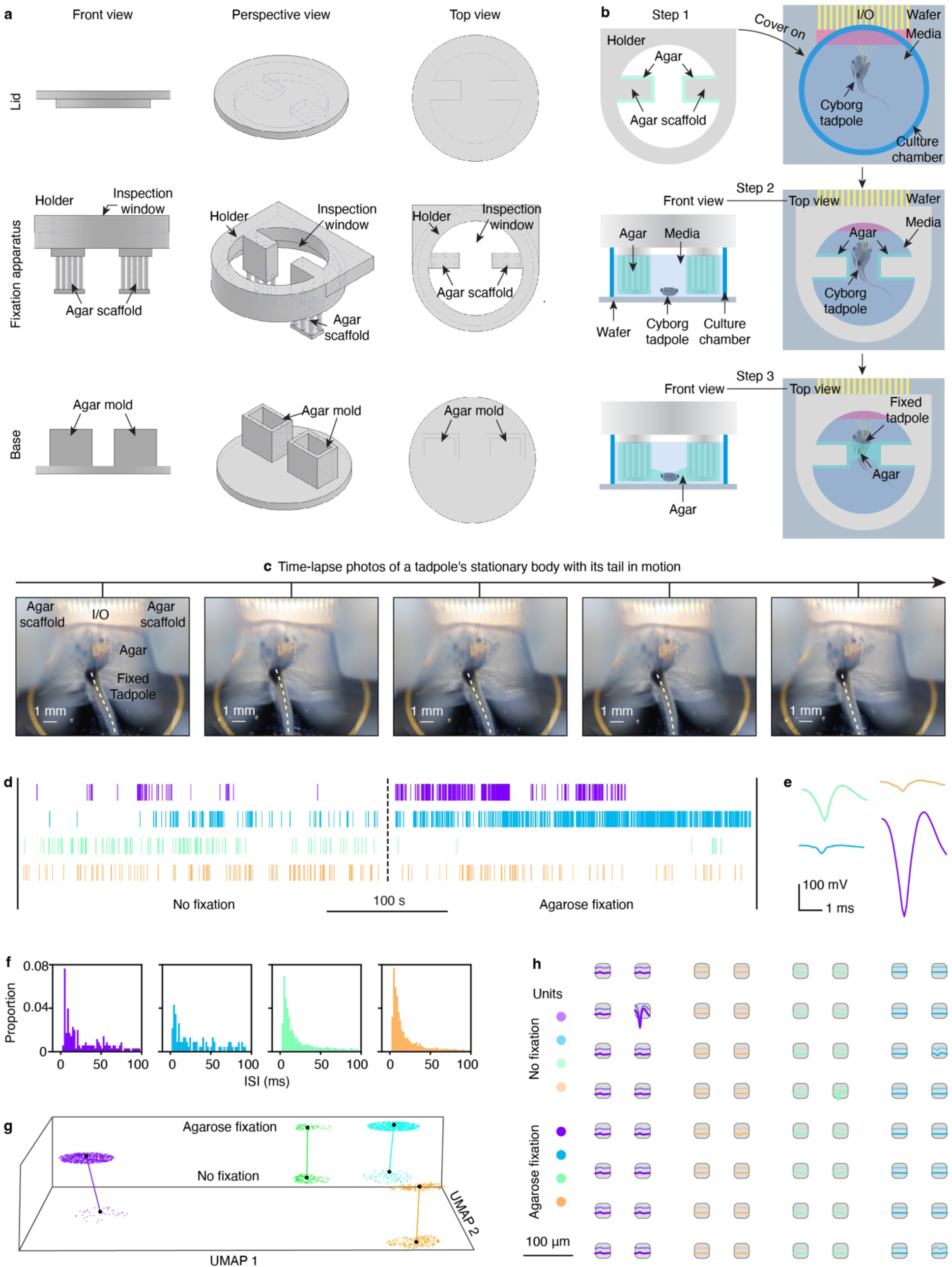
Neural recording in awake cyborg tadpole. **a,** Front, perspective, and top views of the 3D-printed holder for the head-fixed recording. **b,** Schematics showing the stepwise fixation of the cyborg tadpole for awake recording. Step 1: A layer of agarose is cured on the agar scaffold of the holder. Step 2: The holder is placed on top of the culture chamber containing the anesthetized cyborg tadpole. The design of the holder ensures that the tadpole fits in the middle of the two agarose scaffolds. Step 3: A small amount of low melting point agarose further fixes the tadpole with the scaffolds. **c,** Time-lapse photographic images showing that in this setting, the tail of the agarose-fixed tadpole can still move during recording. **d,** Raster plot of spikes sorted from a representative recording of a cyborg tadpole with and without agarose fixation. **e, f,** Average waveforms (**e**) and ISI (**f**) of spikes sorted from recordings in (**d**). **g,** Uniform manifold approximation and projection (UMAP) analysis of neurons from recordings with and without agarose fixation. **h,** Representative average single-unit waveforms at each of the electrodes from recordings with and without agarose fixation. **d-h,** Each color corresponds to an identified unit.

